# Recessive genetic effects on type 2 diabetes-related metabolites in a consanguineous population

**DOI:** 10.1101/619262

**Authors:** Ayşe Demirkan, Jun Liu, Najaf Amin, Jan B van Klinken, Ko Willems van Dijk, Cornelia M. van Duijn

**Affiliations:** Genetic Epidemiology Subunit, Department of Epidemiology, Erasmus University Medical Center, Rotterdam, the Netherlands; Department of Experimental and Clinical Research, Faculty of Bioscience and Medicine, University of Surrey, Guildford, UK; Department of Genetics, University Medical Center Groningen, Groningen, the Netherlands; Nuffield Department of Population Health, University of Oxford, Oxford, UK; Department of Human Genetics, Leiden University Medical Center, Leiden, the Netherlands; Department of Endocrinology, Leiden University Medical Center, Leiden, the Netherlands; Leiden Academic Center for Drug Research, Leiden University, Leiden, the Netherlands

**Keywords:** Type 2 diabetes, metabolomics, homozygosity, autozygosity

## Abstract

Autozygosity, meaning inheritance of an ancestral allele in the homozygous state is known to lead bi-allelic mutations that manifest their effects through the autosomal recessive inheritance pattern. Autosomal recessive mutations are known to be the underlying cause of several Mendelian metabolic diseases, especially among the offspring of related individuals. In line with this, inbreeding coefficient of an individual as a measure of cryptic autozygosity among the general population is known to lead adverse metabolic outcomes including Type 2 diabetes (T2DM); a multifactorial metabolic disease for which the recessive genetic causes remain unknown. In order to unravel such effects for multiple metabolic facades of the disease, we investigated the relationship between the excess of homozygosity and the metabolic signature of T2DM. We included a set of 53 metabolic phenotypes, including 47 metabolites, T2DM and five T2DM risk factors, measured in a Dutch genetic isolate of 2,580 people. For 20 of these markers, we identified 29 regions of homozygous (ROHs) associated with the nominal significance of P-value < 1.0 × 10^−3^. By performing association according to the recessive genetic model within these selected regions, we identified and replicated two intronic variants: rs6759814 located in *KCNH7* associated with valine and rs1573707 located in *PTPRT* associated with IDL-free cholesterol and IDL-phospholipids. Additionally, we identified a rare intronic SNV in *TBR1* for which the homozygous individuals were enriched for obesity. Interestingly, all three genes are mainly neuronally expressed and pointed out the involvement of glutamergic synaptic transmission pathways in the regulation of metabolic pathways. Taken together our study underline the additional benefits of model supervised analysis, but also seconds the involvement of the central nervous system in T2DM and obesity pathogenesis.

## Introduction

Consanguineous marriages between close relatives as a result of assertive mating is known to cause severe metabolic diseases in the off-spring(Vernon 2015). In addition to that, moderate inbreeding due to isolation in populations has been shown to cause unfavorable outcomes among with cardio-metabolic and neuropsychiatric parameters(Verweij *et al*. 2014; Howrigan *et al*. 2016). The previous report shows evidence that inbreeding associates with an increase in blood pressure, glucose and decrease in high-density lipoprotein cholesterol (HDL-C), intelligence quotient (IQ) and height(Mcquillan *et al*. 2012). In the decade, technological advances in metabolomics allow researchers to capture the biochemical status in an organism. As one of the most common metabolic disorders, studying the metabolomics of type 2 diabetes (T2DM) is particularly promising as the deregulation of biochemical processes is involved in the pathophysiology of T2DM. In line with this, many circulating metabolites have been found associated with T2DM: including phospholipids, branch-chain amino-acids and lipoprotein subclasses(Wang *et al*. 2005; Wang *et al*. 2011; Liu *et al*. 2017).

In our previous report, we tested the evidence of recessive SNP effects. We revealed that six out of the eight quantitative metabolite genetic loci showed a recessive rather than an additive effect on the metabolite (Demirkan *et al*. 2015). This raises an important question of whether such recessive variants are relevant for metabolic diseases such as T2DM and its markers. In order to answer this question and find genetic loci that act under recessive inheritance, we first defined runs of homozygosity (ROHs) that relate to T2DM and its circulating profile in blood in the genetically isolated ERF population and then looked for the causal recessive variants within the loci using coding variants. We focused on a set of 47 metabolites which we selected based on their correlation to inbreeding in the population(Demirkan *et al*. 2019). On top of that, we additionally studied the five commonly measured metabolic phenotypes, i.e. body-mass index (BMI), waist-hip ratio (WHR), fasting glucose, insulin and homeostatic model assessment for insulin resistance (HOMA-IR), as well as T2DM case/control status in the ERF population.

## Methods

### Study population

The Erasmus Rucphen Family genetic isolate study (ERF) is a prospective family-based study located in Southwest of the Netherlands. This young genetic isolate was founded in the mid-eighteenth century and minimal immigration and marriages occurred between surrounding settlements due to social and religious reasons. The study population includes 3,465 individuals that are living descendants of 22 couples with at least six children baptized. Informed consent has been obtained from patients where appropriate. The study protocol was approved by the medical ethics board of the Erasmus Medical Center Rotterdam, the Netherlands(Santos *et al*. 2006). The baseline demographic data and measurements of the ERF participants were collected around 2002 to 2006. All the participants filled out questionnaires on socio-demographics, diseases and medical history and lifestyle factors, and were invited to the research center for an interview and blood collection for biochemistry and physical examinations including blood pressure and anthropometric measurements have been performed. The participants were asked to bring all their current medications for registration during the interview. Venous blood samples were collected after at least eight hours of fasting. Baseline type 2 diabetes was defined according to the fasting plasma glucose ≥ 7.0mmol/L and/or anti-diabetic treatment, yielding 212 cases and 2,564 controls, totaling up to 2,776. The follow-up data collection of the ERF study took place in May 2016 (9 to 14 years after baseline visit). During the follow up a total of 1,935 participants’ records were scanned for the incidence of type 2 diabetes in general practitioner’s databases. Additionally, a questionnaire on type 2 diabetes medication surveyed on 1,232 participants in June 2010 (4 to 8 years after baseline visit) was referred if a participant were not included in May 2016 follow-up. This effort yielded the inclusion of 18 otherwise missed extra cases, yielding a total of 349 cases and 2,427 controls in the current study.

### Metabolite measurements

Metabolic markers were measured by five different metabolomics platforms using the methods which have been described in earlier publications(Demirkan *et al*. 2012; Gonzalez-Covarrubias *et al*. 2013; Demirkan et al. 2015; Draisma *et al*. 2015; Kettunen *et al*. 2016). In total 562 metabolic markers including sub-fractions of lipoproteins, triglycerides, phospholipids, ceramides, amino acids, acyl-carnitines and small intermediate compounds, which throughout this article will be referred as “*metabolites*”, were measured either by nuclear magnetic resonance (NMR) spectrometry or by mass spectrometry (MS). The platforms used in this research are: (1) Liquid Chromatography-MS (LC-MS, 116 positively charged lipids, comprising of 39 triglycerides, 47 phosphatidylcholines, 8 phosphatidylethanolamines, 20 sphingolipids, and 2 ceramides, available in up to 2,638 participants) measured in the Netherlands Metabolomics Center, Leiden using the method described before(Gonzalez-Covarrubias *et al*. 2013); (2) Electrospray-Ionization MS (ESI-MS, in total 148 phospholipids and sphingolipids comprising of 16 plasmologens, 72 phosphatidylcholines, 27 phosphatidylethanolamines, 24 sphingolipids, 9 ceramides, available in up to 878 participants), measured in the Institute for Clinical Chemistry and Laboratory Medicine, University Hospital Regensburg, Germany using the method described previously (Demirkan *et al*. 2012); (3) Small molecular compounds window based NMR spectroscopy (41 molecules comprising of 29 low-molecular weight molecules and 12 amino acids available in up to 2,639 participants) measured in the Center for Proteomics and Metabolomics, Leiden University Medical Center(Demirkan *et al*. 2015; Verhoeven *et al*. 2017); (4) Lipoprotein window-based NMR spectroscopy (104 lipoprotein particles size sub-fractions comprising of 28 VLDL components, 30 HDL components, 35 LDL components, 5 IDL components and 6 plasma totals, available in 2,609 participants) measured in the Center for Proteomics and Metabolomics, Leiden University Medical Center and lipoprotein sub-fraction concentrations were determined by the Bruker algorithm (Bruker BioSpin GmbH, Germany) as detailed in Kettunen *et al*(Kettunen *et al*. 2016); (5) AbsoluteIDQTM p150 Kit of Biocrates Life Sciences AG (153 molecules comprising of 14 amino acids, 91 phospholipids, 14 sphingolipids, 33 acyl-carnitines and hexose available in up to 989 participants) measured as detailed in publication from Draisma *et al*(Draisma *et al*. 2015) and the experiments were carried out at the Metabolomics Platform of the Genome Analysis Center at the Helmholtz Zentrum München, Germany as per the manufacturer’s instructions. The laboratories had no access to phenotype information.

### Genome-wide SNP measurements

Genotyping in ERF was performed using Illumina 318/370 K, Affymetrix 250 K, and Illumina 6K micro-arrays. All SNPs were imputed using MACH software (www.sph.umich.edu/csg/abecasis/MaCH/) based on the Hapmap2 (release 22, build 36). Individuals were excluded for excess autosomal heterozygosity, mismatches between called and phenotypic gender, and if there were outliers identified by an identical-by-state (IBS) clustering analysis.

### Defining ROHs and regression

We used Hapmap2 (release 22, build 36) imputations for the extraction of homozygosity per loci per individual. The best guess information used for SNPs with imputation quality (R^2^) > 0.95. The algorithm used to define the region-wise homozygosity was developed by the ROHGEN consortium as explained before(Ceballos *et al*. 2018). In brief, the genome was divided into 3 Mb windows (n=992) and for each window, a plink sliding window routine was used to identify the proportion of >1.5 Mb homozygosity within the window. For each window, a maximum of one heterozygous SNP and five missing measurements were allowed. All SNP ids were mapped to the human genome build 19 (hg19) coordinates. The regression analyses were performed using mixed models, adjusting for genetic relatedness using a genomic kinship matrix; age, sex, and first 10 principal components were used as covariates in the model.

### Exome sequence measurements

Exomes of 1,336 randomly selected individuals from the ERF study cohort were sequenced at the Center for Biomics of the Cell Biology department, at the Erasmus MC, The Netherlands. Sequencing was done at a median depth of 57× using the Agilent version V4 capture kit, on an Illumina Hiseq2000 sequencer, using the TruSeq Version 3 protocol. The sequence reads were aligned to hg19, using Burrows-Wheeler Aligner (BWA) and the NARWHAL pipeline(Li and Durbin 2009), (Brouwer *et al*. 2012) (Li *et al*. 2009). Subsequently, the aligned reads were processed further, using the IndelRealigner, MarkDuplicates and Table Recalibration tools from the Genome Analysis Toolkit (GATK)(Mckenna *et al*. 2010) and Picard (http://picard.sourceforge.net). This was necessary to remove systematic biases and to recalibrate the PHRED quality scores in the alignments. After processing, genetic variants were called, using the Unified Genotyper tool from the GATK(Mckenna *et al*. 2010). For each sample, at least four Gigabases of sequence was aligned to the genome. Functional annotations were also performed using the SeattleSeq annotation 138 database. About 1.4 million SNVs were called. After removing variants with low quality, out of Hardy-Weinberg equilibrium (HWE, P-value < 10^−6^) and low call rate (< 99%), and samples with a low call rate (< 90%), we retrieved 543,954 very high-quality SNVs in 1,327 individuals.

### Exome chip measurements

Study participants from the ERF study whose exomes were not sequenced (N = 1,527) were genotyped on the Illumina Infinium HumanExome BeadChip, version 1.1, which contains over 240,000 exonic variants selected from multiple sources together spanning 12,000 samples from multiple ethnicities. Calling was performed with GenomeStudio. We removed subjects with a call rate < 95%, IBS > 0.99 and heterozygote ratio > 0.60. Ethnic outliers identified using a principal component analysis with 1000 Genomes data and individuals with sex discrepancies were also removed. The SNVs that were monomorphic in our sample or had a call rate < 95% were removed. After quality control, we retrieved about 70,000 polymorphic SNVs in 1,515 subjects.

### Genetic association analyses

#### Exome chip and sequence-based genetic variants

All tests were performed using RVtests (version 20150630), adjusted by age, sex and familial relatedness using a genome wide-autosomal kinship matrix(Zhan *et al*. 2016). Single variant association analyses were performed for variants that have more than five copies of homozygous presentation in the population, using a recessive genetic model.

#### Imputation-based genetic variants

The SNPs with less than five minor allele counts were excluded. The exclusion criteria for SNPs were HWE with P-value < 1.0 × 10^−6^ or SNP call rate < 98%. The associations of the significant metabolites within the genotype in these regions were performed by linear regression with ProbABEL software in both recessive model and dominant model in order to capture the homozygous model for both effect and non-effect alleles. The residuals of the metabolites with age, sex and family-relationship matrix were calculated by the *polygenic* function in R package GenABEL and used in the region-wide associations. The associations within the regions were performed using the 1000G phase 1, release v3. All SNP ids were mapped to hg19 coordinates. The genotype data of a population-based cohort, Rotterdam Study(Ikram *et al*. 2017) (n = 6,291), was also used to replicate the findings.

Candidate genes within the ROH were selected by in-house developed automated pathway search algorithm as explained before(Demirkan *et al*. 2015). The selection algorithm combines information from GTEx-eQTL, GWAS catalog, ConsensusPathDB(Kamburov *et al*. 2011), UniProtKB(Magrane 2011), OMIM(Mckusick 1998), TCDB(Saier JR *et al*. 2006), ExPASy(Gasteiger *et al*. 2003) and KEGG database(Kanehisa and Goto 2000) for each genetic loci.

## Results

### ROH regression analysis

In total, 47 metabolites selected in ERF population in an earlier report(Demirkan *et al*. 2019) in addition to T2DM and five T2DM-related phenotypes were included in the ROH regression analysis with 992 windows constructed genome-wide. These analyses yielded 3,334 window-metabolite pairs with association p-value < 0.05. However, none of the associations passed either the false discovery rate (FDR), or Bonferroni threshold defined as correction by the number of windows tested times the independent dimensions in the highly correlated data(Li and Ji 2005) (0.05/ (992 × 27) = 2.02 × 10^−6^). Out of 3,334, 1,734 association pairs showed consistent direction with the initially calculated association to inbreeding coefficients from our earlier report(Demirkan *et al*. 2019). We focused on the suggestive top loci with association P-value < 1.0 × 10^−3^ for further analysis. These associations with ROH and phenotypes are given in Table 1. The list included 29 genomic loci identified as full or partial homozygosity within a 3MB, influencing 20 different outcomes, in total making up 51 suggestive pairs of association for follow-up. In some loci, the neighboring regions were correlated indicating that the ROH detected was indeed larger than 3MB window **(Supplementary Figure 1).** The top significant ROH which is shared by 33 individuals is located at 36 to 39 MB on chromosome 4 and is associated with four small size LDL components (S-LDL-Free cholesterol, S-LDL-phospholipids, S-LDL-cholesterol, S-LDL-ApoB) and included candidate genes *TLR* and *PGM2*. The second top ROH shared by 43 individuals is located at 63 to 66 MB on chromosome 3, and is associated with a lifetime risk of T2DM (β = 0.43, P-value=5.43 × 10^−6^) and included candidate genes *PSMD6* and *ADAMTS9* (Table 1). ROH located at chromosome 4, 30-33 Mb was highest frequently shared (420 individuals). The strongest association was found between this ROH and S-LDL-Free cholesterol (β = 1.35, P-value = 7.48 × 10^−5^), followed by S-LDL-cholesterol, S-LDL-phospholipids and S-VLDL-triglycerides. For the rest 26 top ROH the number of carrier individuals ranged between 20 to 118. Distributions of the 29 ROH in ERF population are provided in **Supplementary Figure 2**.

**Table 1.**
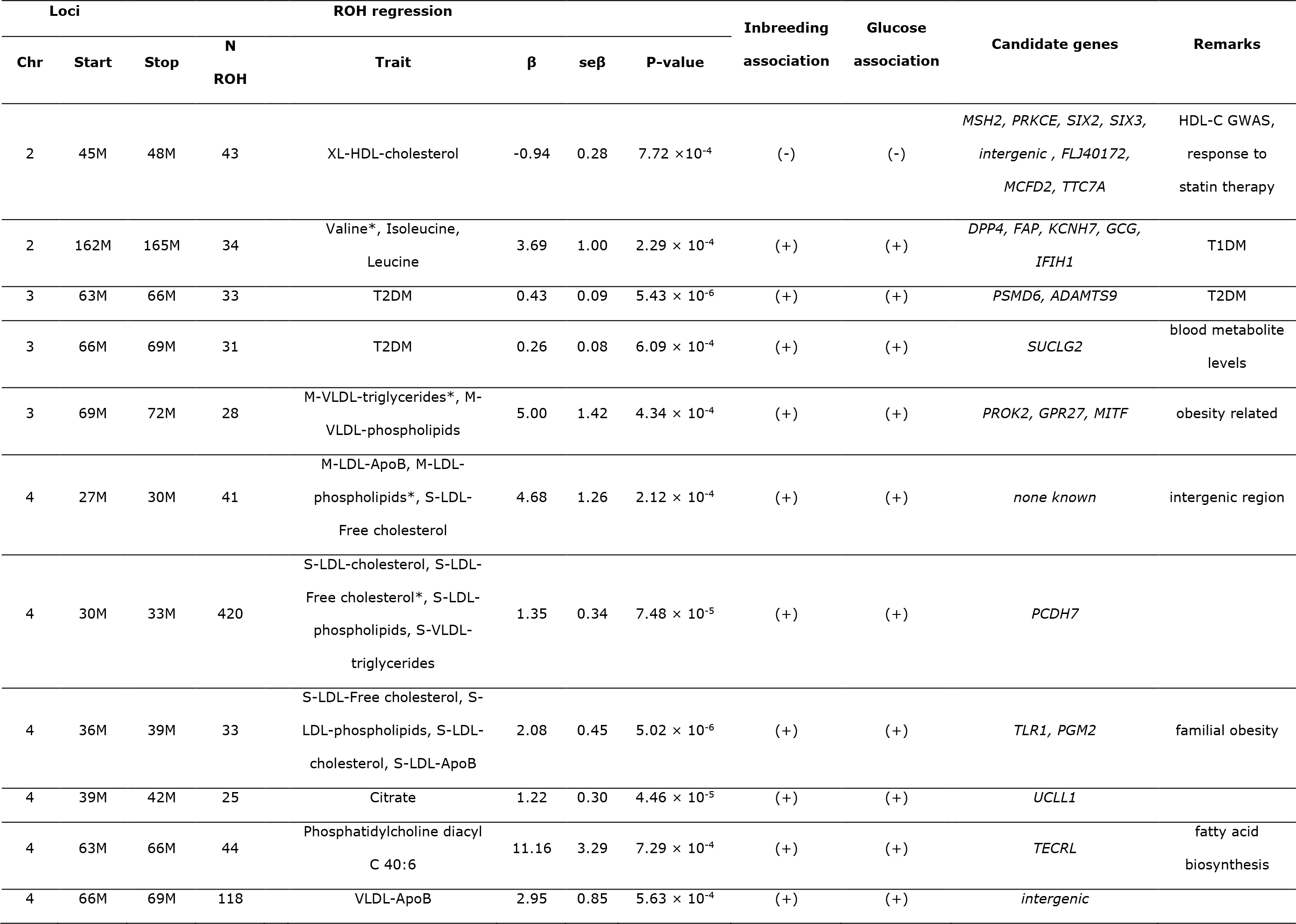
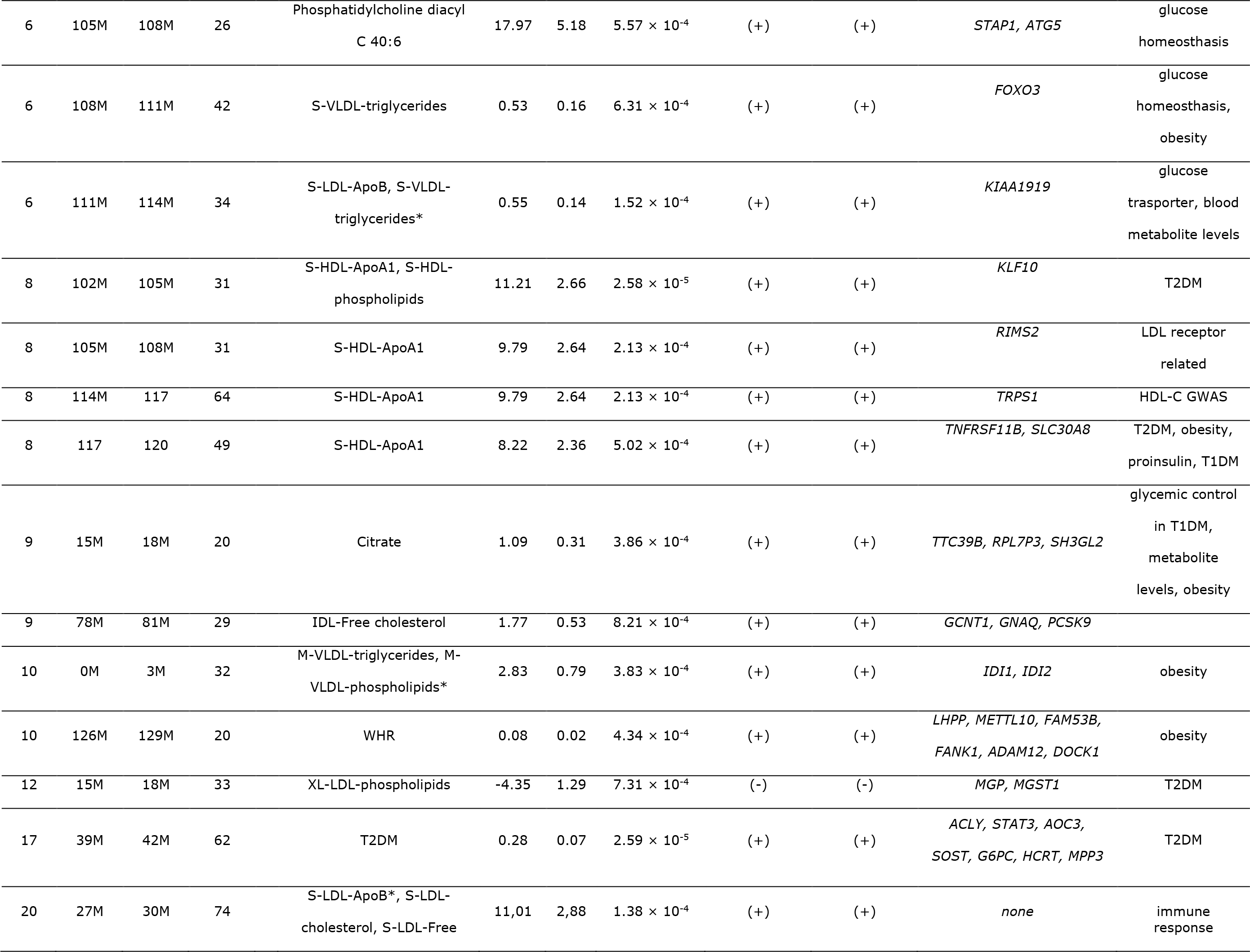
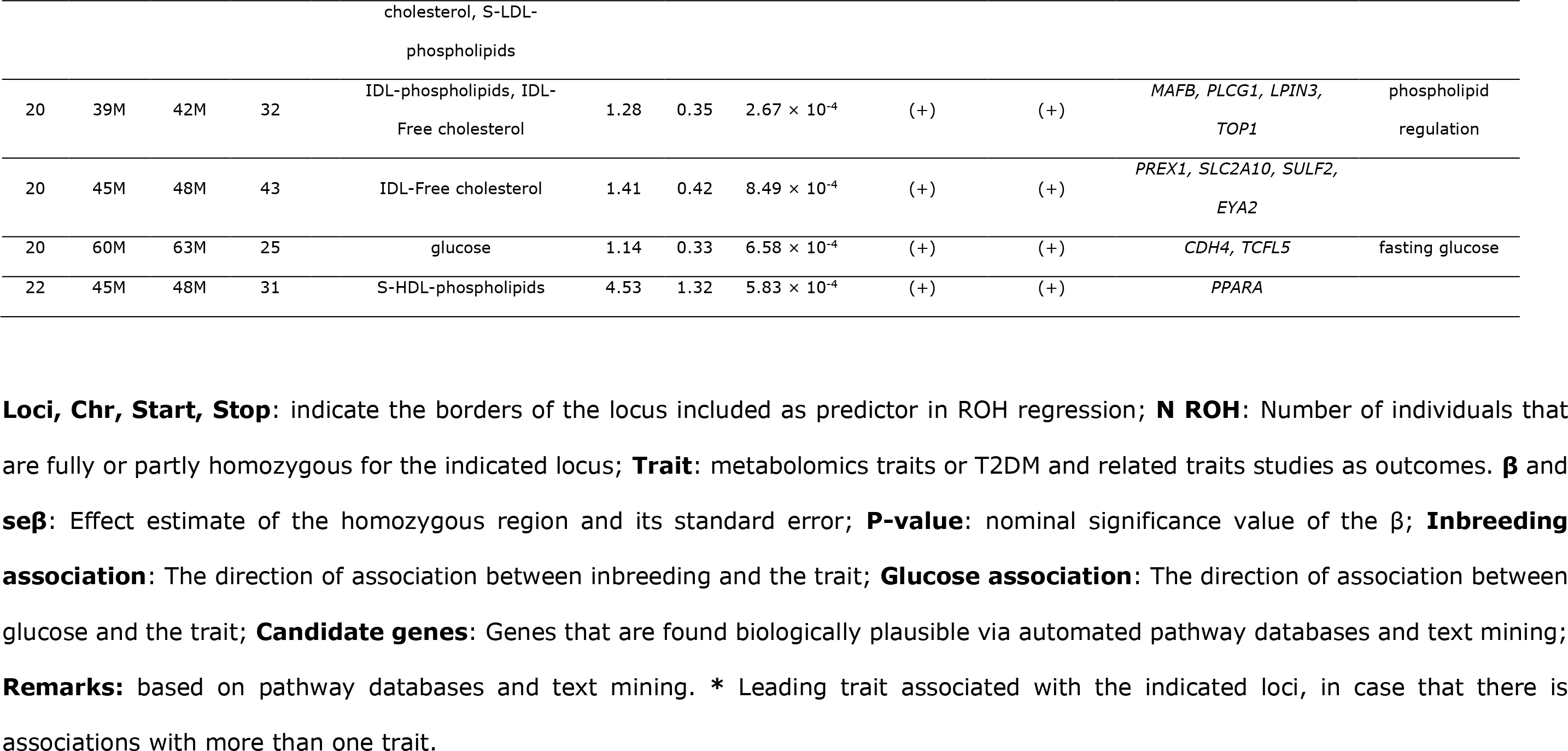
ROH regression homozygosity mapping top results with P-value < 1.0 × 10^−3^

### Fine mapping within the ROHs

#### Association with exonic variants

For each locus, the association for the minor allele under the recessive models has been tested, results are shown in Table 2a and Table 2b with sequence and chip-based genotyping sets respectively. We used three steps of variant filtering. First, we adopted a liberal variant list and included all the SNVs that are captured within the sequence data, also including the intergenic and intronic variants that are captured around the exons. For each region, we set up a region-wide Bonferroni threshold based on the number of SNVs tested and based on that we identified the significantly associated SNVs. By this way five SNVs from exonic sequences were detected inside genes *TTC7A (*rs57182920, intronic, with XL-HDL-Cholesterol*), FRMD4B (*rs73095903, intronic, with M-VLDL-TGs*), CSMD3* (rs72685825, intronic, with S-HDL-ApoA1), *PREX1 (*rs3746820, synonymous, with IDL-phospholipids*), LAMA5 (*rs35653162, intronic, with glucose) in addition to one SNV detected inside a non-transcribed pseudogene *EXOC5P1* (rs6551721, exonic, with Phosphatidylcholine diacyl C 40:6) as shown in Table 2 and **Supplementary Table 1**. By using the exome chip derived genetic data, we detected three SNVs; one located inside *KCNH7 (*rs6759814, intronic, with valine), and two from intergenic regions (rs6469084, associated with S-HDL-ApoA1 and rs6101991 associated with IDL-phospholipids) detected passing the pre-defined region-specific association thresholds (Table 3 and **Supplementary Table 2**). Second, we limited the analysis to synonymous and non-synonymous SNVs stop codons, UTR and splice variants only. By this way, additional two synonymous SNVs located in *OXR1* (rs1681904, with S-HDL-ApoA1) and *FRMD4B* (rs62254461, with M-VLDL-triglycerides) were found using the exonic sequences (Table 2). Using the same approach one missense coding SNV in the exome chip was found associated within *KRT15* gene (rs1050784, with T2DM) (Table 3). Third, we filtered the dataset such that we focused on only to those missense and premature stop codons. However, no SNVs were found associated using such strict filtering.

**Table 2.**
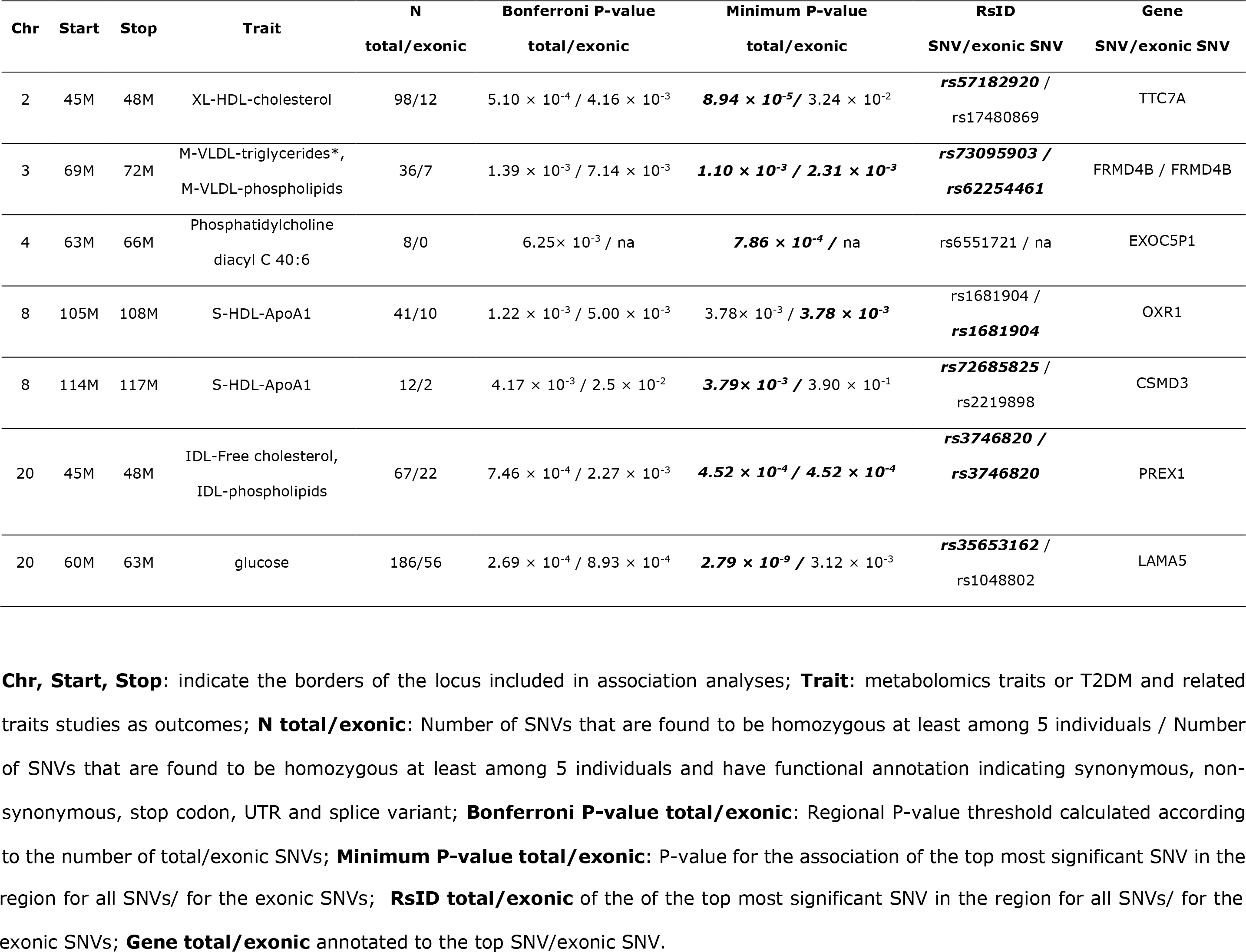
Association with exonic SNVs in the loci (NGS based measurements) using the recessive genetic model

| Chr | Start | Stop | Trait | N<br>total/exonic | Bonferroni P-value<br>total/exonic | Minimum P-value<br>total/exonic | RsID<br>SNV/exonic SNV | Gene<br>SNV/exonic SNV |
| --- | --- | --- | --- | --- | --- | --- | --- | --- |
| 2 | 45M | 48M | XL-HDL-cholesterol | 98/12 | $5.10 \times 10^{-4} / 4.16 \times 10^{-3}$ | $8.94 \times 10^{-5} / 3.24 \times 10^{-2}$ | **rs57182920** /<br>rs17480869 | TTC7A |
| 3 | 69M | 72M | M-VLDL-triglycerides*,<br>M-VLDL-phospholipids | 36/7 | $1.39 \times 10^{-3} / 7.14 \times 10^{-3}$ | $1.10 \times 10^{-3} / 2.31 \times 10^{-3}$ | **rs73095903** /<br>**rs62254461** | FRMD4B / FRMD4B |
| 4 | 63M | 66M | Phosphatidylcholine<br>diacyl C 40:6 | 8/0 | $6.25 \times 10^{-3} / \text{na}$ | $7.86 \times 10^{-4} / \text{na}$ | rs6551721 / na | EXOC5P1 |
| 8 | 105M | 108M | S-HDL-ApoA1 | 41/10 | $1.22 \times 10^{-3} / 5.00 \times 10^{-3}$ | $3.78 \times 10^{-3} / 3.78 \times 10^{-3}$ | rs1681904 /<br>**rs1681904** | OXR1 |
| 8 | 114M | 117M | S-HDL-ApoA1 | 12/2 | $4.17 \times 10^{-3} / 2.5 \times 10^{-2}$ | $3.79 \times 10^{-3} / 3.90 \times 10^{-1}$ | **rs72685825** /<br>rs2219898 | CSMD3 |
| 20 | 45M | 48M | IDL-Free cholesterol,<br>IDL-phospholipids | 67/22 | $7.46 \times 10^{-4} / 2.27 \times 10^{-3}$ | $4.52 \times 10^{-4} / 4.52 \times 10^{-4}$ | **rs3746820** /<br>**rs3746820** | PREX1 |
| 20 | 60M | 63M | glucose | 186/56 | $2.69 \times 10^{-4} / 8.93 \times 10^{-4}$ | $2.79 \times 10^{-9} / 3.12 \times 10^{-3}$ | **rs35653162** /<br>rs1048802 | LAMA5 |

**Chr, Start, Stop:** indicate the borders of the locus included in association analyses; **Trait:** metabolomics traits or T2DM and related traits studies as outcomes; **N total/exonic:** Number of SNVs that are found to be homozygous at least among 5 individuals / Number of SNVs that are found to be homozygous at least among 5 individuals and have functional annotation indicating synonymous, non-synonymous, stop codon, UTR and splice variant; **Bonferroni P-value total/exonic:** Regional P-value threshold calculated according to the number of total/exonic SNVs; **Minimum P-value total/exonic:** P-value for the association of the top most significant SNV in the
region for all SNVs/ for the exonic SNVs; **RsID total/exonic** of the of the top most significant SNV in the region for all SNVs/ for the exonic SNVs; **Gene total/exonic** annotated to the top SNV/exonic SNV.

**Table 3.**
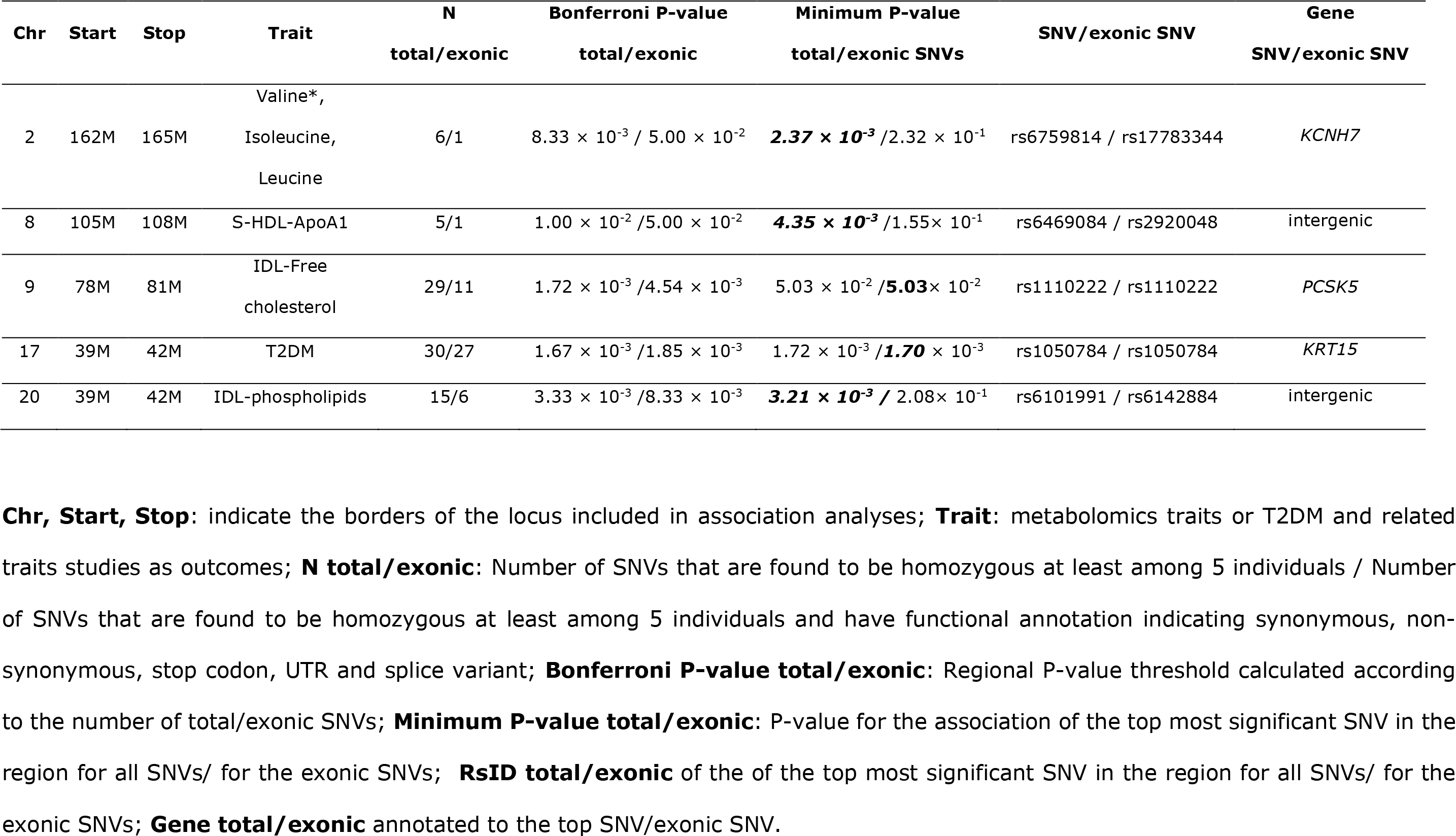
Association with exonic SNVs in the loci (chip based measurements) using the recessive genetic model

| Chr | Start | Stop | Trait | N<br>total/exonic | Bonferroni P-value<br>total/exonic | Minimum P-value<br>total/exonic SNVs | SNV/exonic SNV | Gene<br>SNV/exonic SNV |
| --- | --- | --- | --- | --- | --- | --- | --- | --- |
| 2 | 162M | 165M | Valine*,<br>Isoleucine,<br>Leucine | 6/1 | $8.33 \times 10^{-3} / 5.00 \times 10^{-2}$ | **$2.37 \times 10^{-3}$** / $2.32 \times 10^{-1}$ | rs6759814 / rs17783344 | *KCNH7* |
| 8 | 105M | 108M | S-HDL-ApoA1 | 5/1 | $1.00 \times 10^{-2} / 5.00 \times 10^{-2}$ | **$4.35 \times 10^{-3}$** / $1.55 \times 10^{-1}$ | rs6469084 / rs2920048 | intergenic |
| 9 | 78M | 81M | IDL-Free<br>cholesterol | 29/11 | $1.72 \times 10^{-3} / 4.54 \times 10^{-3}$ | $5.03 \times 10^{-2} /$**$5.03 \times 10^{-2}$** | rs1110222 / rs1110222 | *PCSK5* |
| 17 | 39M | 42M | T2DM | 30/27 | $1.67 \times 10^{-3} / 1.85 \times 10^{-3}$ | $1.72 \times 10^{-3} /$**$1.70 \times 10^{-3}$** | rs1050784 / rs1050784 | *KRT15* |
| 20 | 39M | 42M | IDL-phospholipids | 15/6 | $3.33 \times 10^{-3} / 8.33 \times 10^{-3}$ | **$3.21 \times 10^{-3}$** / $2.08 \times 10^{-1}$ | rs6101991 / rs6142884 | intergenic |

**Chr, Start, Stop:** indicate the borders of the locus included in association analyses; **Trait:** metabolomics traits or T2DM and related traits studies as outcomes; **N total/exonic:** Number of SNVs that are found to be homozygous at least among 5 individuals / Number of SNVs that are found to be homozygous at least among 5 individuals and have functional annotation indicating synonymous, non-synonymous, stop codon, UTR and splice variant; **Bonferroni P-value total/exonic:** Regional P-value threshold calculated according to the number of total/exonic SNVs; **Minimum P-value total/exonic:** P-value for the association of the top most significant SNV in the region for all SNVs/ for the exonic SNVs; **RsID total/exonic** of the of the top most significant SNV in the region for all SNVs/ for the exonic SNVs; **Gene total/exonic** annotated to the top SNV/exonic SNV.

#### Association with common variants and intergenic regions

Table 4 and **Supplementary Table 3** shows the region-wide association with 1000 genome imputed genotypes within the candidate ROHs. The locus zoom plots of loci with significant SNPs are given in **Supplementary Figure 3.** In total, common genetic variants in six regions passed the region-wide FDR. These included rs59997916 (intronic *MAGI1*, with T2DM), rs10866392 (intronic LINCO2506, with S-LDL-Free cholesterol), rs73240383 (intronic, *NWD2*, with S-LDL-ApoB), rs71562230 (intergenic near *SLC22A16*, with S-VLDL-triglycerides), rs1573707 (intronic, *PTPRT*, with IDL-Free cholesterol, IDL-phospholipids) and rs75320186 (upstream of *BIRC7*, with glucose).

**Table 4.**
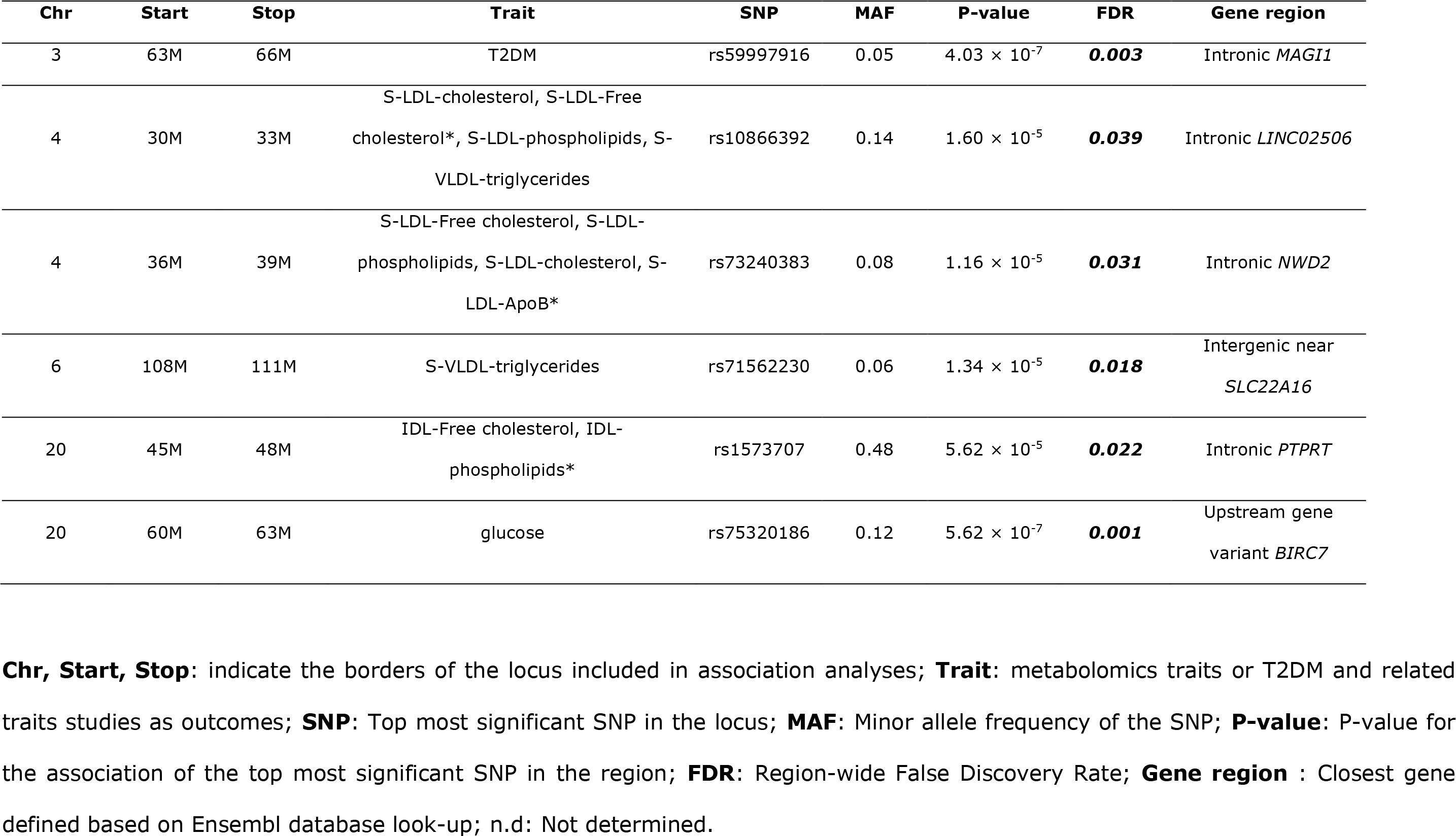
Association analyses of imputed SNPs within the candidate ROHs using the recessive genetic model

In order to investigate the effect of these variants in the outbred population, Rotterdam Study we set out a replication. Out of the 18 genetic associations coming either from exome sequencing, exome chip or imputation sets, we were able to test 12 in the Rotterdam study as 6 of the phenotypes were not available for replication. (Table 5 and **Supplementary Table 4**). After correcting for multiple testing, we replicated two intronic genetic variants are located in genes *KCNH7* (rs6759814) and *PTPRT* (rs1573707).

**Table 5.**
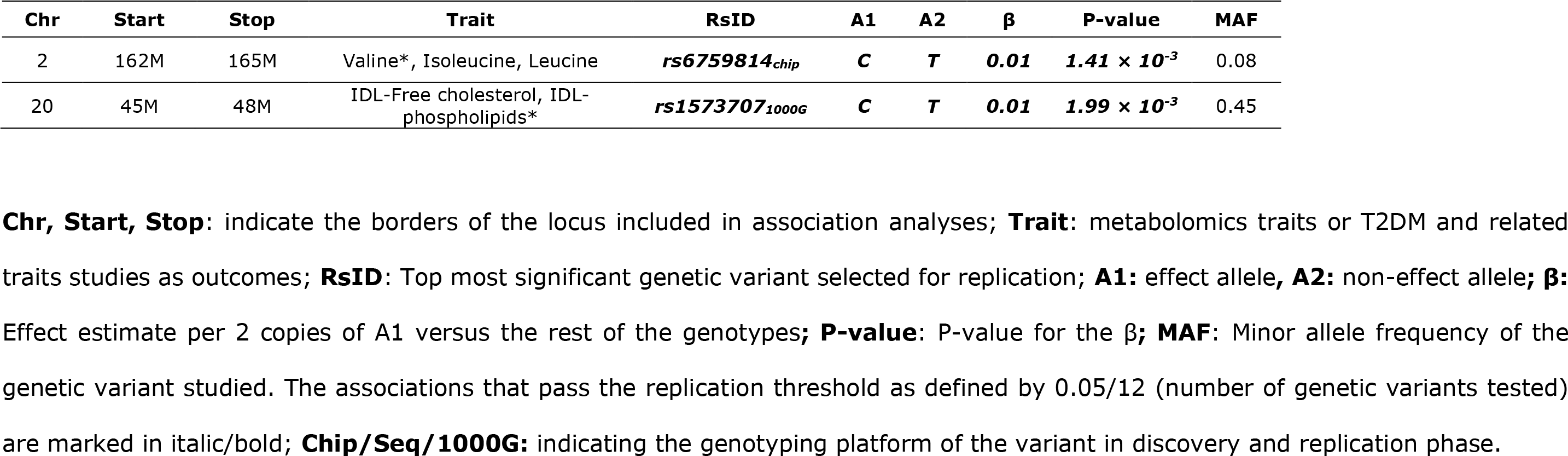
Replication of selected SNVs in Rotterdam Study using the recessive genetic model

| **Chr** | **Start** | **Stop** | **Trait** | **RsID** | **A1** | **A2** | **β** | **P-value** | **MAF** |
| --- | --- | --- | --- | --- | --- | --- | --- | --- | --- |
| 2 | 162M | 165M | Valine*, Isoleucine, Leucine | ***rs6759814<sub>chip</sub>*** | ***C*** | ***T*** | ***0.01*** | ***1.41 × 10<sup>-3</sup>*** | 0.08 |
| 20 | 45M | 48M | IDL-Free cholesterol, IDL-phospholipids* | ***rs1573707<sub>1000G</sub>*** | ***C*** | ***T*** | ***0.01*** | ***1.99 × 10<sup>-3</sup>*** | 0.45 |

**Chr, Start, Stop:** indicate the borders of the locus included in association analyses; **Trait:** metabolomics traits or T2DM and related traits studies as outcomes; **RsID:** Top most significant genetic variant selected for replication; **A1:** effect allele, **A2:** non-effect allele; **β:** Effect estimate per 2 copies of A1 versus the rest of the genotypes; **P-value:** P-value for the β; **MAF:** Minor allele frequency of the genetic variant studied. The associations that pass the replication threshold as defined by 0.05/12 (number of genetic variants tested) are marked in italic/bold; **Chip/Seq/1000G:** indicating the genotyping platform of the variant in discovery and replication phase.

### Shared rare homozygous SNVs within the ROH regions

In parallel to the association analysis explained above, we searched for overweight (BMI > 25.0 kg/m^2^), obese (BMI > 30.0 kg/m^2^) and T2DM cases among the ROH carriers who also share the same rare (MAF <0.05) genetic variants identified by exome sequence or exome chip. By this way, we found additional 28 rare variants were shared exclusively by the ROH carriers within the genomic regions of interest. After exploring the phenotypes of the individuals homozygous for these exclusive 28 rare SNVs, for 18 of them, we detected an enrichment of cases of T2DM and obesity, clustering among 32 homozygous individuals. Eight out of 32 had T2DM (25.0%), 19 were overweight (59.4%) and 11 were obese (34.4%). Among the 18 rare SNVs, rs116175783 located in *TBR1* can also be detected in the Rotterdam Study in eight homozygous persons. Six of them (75.0%) were overweight and one was obese in the Rotterdam Study. In ERF one out of the two homozygous individuals is obese and has T2DM, whereas the other homozygous individual is overweight (Table 6).

**Table 6.**
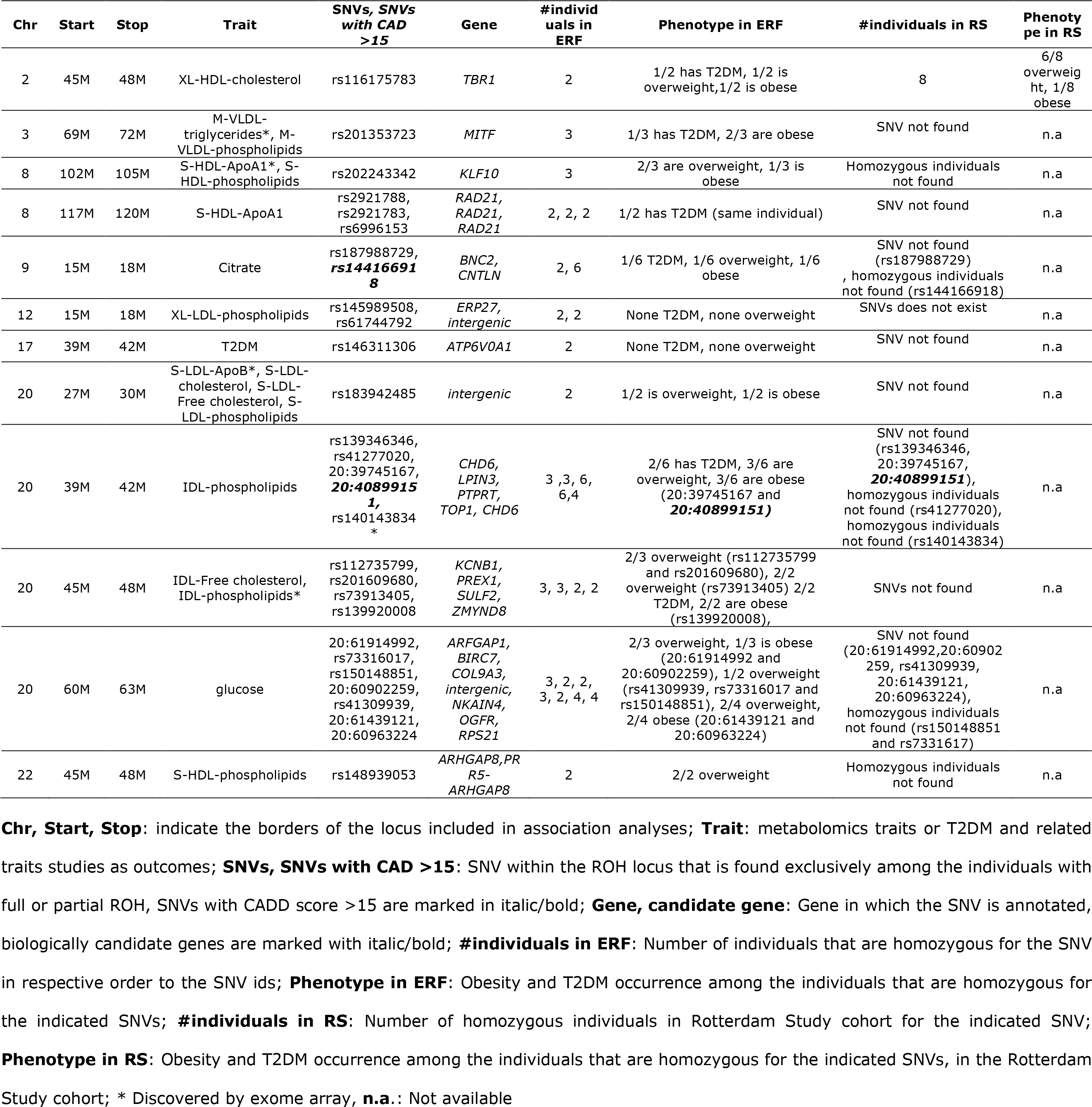
Shared homozygous SNVs specific to the individuals with ROH

## Discussion

We report an in-depth association mapping effort for a total of 53 selected metabolic phenotypes by using three sets of independently generated genotype data. First, we point out increased homozygosity in 29 genomic loci influencing 20 different outcomes, suggesting 51 outcome-genomic locus pairs to be investigated by more in-depth analysis. By using the recessive genetic model for association testing, we detected and replicated 2 intronic genetic variants inside genes *KCNH7* and *PTPRT*. Additionally, within the ROHs we report 18 rare variants in a group of individuals enriched by cases of obesity and T2DM. Among these 18 rare variants, 17 were specific to ERF population, whereas one (rs116175783, located in *TBR1*) was also found in the Rotterdam study, and remarkably six out of eight homozygous individuals are overweight.

Evolutionary selection is less effective in eliminating recessive deleterious alleles since it needs two copies to reduce the fitness of the organism, those tend to become more frequent than expected. The most favorable setting to observe the effect of recessive deleterious genetic variants is consanguineous families or inbred populations. Harmful effects of close consanguinity in humans have been shown for several of outcomes including intelligence(Fareed and Afzal 2014), schizophrenia(Mansour *et al*. 2010), bipolar disorder(Mansour *et al*. 2009), hypertension(Rudan *et al*. 2003), heart disease(Shami *et al*. 1991), cancer(Lebel and Gallagher 1989), but notably also for metabolic health(Isaacs *et al*. 2007) and T2DM(Gosadi *et al*. 2014). Since consanguinity is strongly associated with risk for multifactorial disease, it is more than likely that unknown homozygous regions of the genome explain a significant portion of familial aggregation among other possible mechanisms(Hoppenbrouwers *et al*. 2007). ROHs have been associated with several human traits such as personality(Verweij *et al*. 2012), schizophrenia(Keller *et al*. 2012), short stature(Mcquillan *et al*. 2012) and birth height(Verweij *et al*. 2014).

To our knowledge, this is the first study investigating metabolomics phenotypes by homozygosity mapping. We show that the homozygosity of the minor C allele of synonymous SNV rs6759814 located in *KCNH7* associates with branch chain amino acid valine in our study. *KCNH7* belongs to the ERG subfamily of voltage-gated potassium channels and is widely expressed by the central nervous system. Voltage-gated potassium channels have diverse functions including regulating neurotransmitter release, heart rate, insulin secretion, neuronal excitability, epithelial electrolyte transport, smooth muscle contraction, and cell volume.

Earlier we showed an association of another intronic genetic variant (rs1474260) in *KCNH7* and a polyunsaturated ether-phosphatidylcholine (PCae36:5) in a meta-analysis, using the additive genetic model(Demirkan *et al*. 2012). The second replicated SNV is another synonymous intronic variant (rs1573707) in *PTPRT* associated with IDL-Free cholesterol, IDL-phospholipids*. PTPRT* is a member of the protein tyrosine phosphatase (PTP) family. PTPs are known to be signaling molecules that regulate a variety of cellular processes including cell growth, differentiation, mitotic cycle, and oncogenic transformation. It is of note that both *KCNH7* and *PTPRT* are neuronal proteins associated with glutamate receptor signaling pathway (p-value=5.5 × 10^−4^ for *KCNH7* and p-value= 2.6 × 10^−7^ for *PTPRT* gene expression in blood) as classified by Gene Ontology (GO)-biological process terms, and summarized by genenetwork.nl (accession date: 16.4.2019). Additionally, both proteins map to regulation of insulin secretion (p-value=6.6 × 10^−4^ for *PTPRT* and p-value=3.5 × 10^−4^ for *KCNH7*) among the biological processes classified by GO (www.genenetwork.nl, accession date: 16.04.2019). Finally, by looking at shared homozygous SNVs exclusively by the individuals sharing homozygous regions, we show that a rare genetic variant (rs116175783) shared by a total of 10 individuals in both ERF and Rotterdam Study is interesting as all of the individuals are either overweight or obese. *TBR1* is also a neuronal protein mainly involved in the regulation of glutamergic synaptic transmission (P-value = 3.1 × 10^−7^) associated with behavioral fear response (P-value = 2.7 × 10^−7^). Remarkably, feeding behavior (P-value = 8.1 × 10^−3^) is one of the top GO terms for this gene. Based on GWAS using additive genetic model *TBR1* was found associated with gene-based also associates with sodium in blood (P-value = 3.8 × 10^−10^), educational attainment (P-value = 1.8 × 10^−8^), diagnosed High blood pressure (2.7 × 10^−8^) but also with BMI (6.1 × 10^−6^) to some degree (http://atlas.ctglab.nl/PheWAS, accession date: 16.04.2019). In our study, the region harboring *TBR1* was initially selected because the homozygosity was associated with a decrease in XL-HDL-cholesterol level.

Overall by combining the power of recessive inheritance in a genetically isolated population with a wide range of metabolic pathways, we pointed out several genetic loci that could be of interest for further research. By performing fine mapping within these genetic loci we found and replicated two genetic variants in i.e. *KCNH7*, *PTPRT* involved in glutamergic synaptic pathways by using recessive genetic association models. In addition, we found a rare SNV in another synaptic gene, *TBR1*, for which the homozygous carriers are enriched in obese phenotype. Taken together our study underline the additional benefits of model supervised analysis but also seconds the involvement of the central nervous system in T2DM and obesity pathogenesis.

## Acknowledgments

ERF was supported by the Consortium for Systems Biology (NCSB) both within the framework of the Netherlands Genomics Initiative (NGI)/Netherlands Organization for Scientific Research (NWO). ERF study as a part of EUROSPAN (European Special Populations Research Network) was supported by European Commission FP6 STRP grant number 018947 (LSHG-CT-2006-01947) and also received funding from the European Community’s Seventh Framework Program (FP7/2007-2013)/grant agreement HEALTH-F4-2007-201413 by the European Commission under the program “Quality of Life and Management of the Living Resources” of 5th Framework Program (no. QLG2-CT-2002-01254) as well as FP7 project EUROHEADPAIN (nr 602633). High-throughput analysis of the ERF data was supported by a joint grant from Netherlands Organization for Scientific Research and the Russian Foundation for Basic Research (NWO-RFBR 047.017.043). High throughput metabolomics measurements of the ERF study has been supported by BBMRI-NL (Biobanking and Biomolecular Resources Research Infrastructure Netherlands). Ayse Demirkan is supported by a Veni grant (2015) from ZonMw. Ayse Demirkan, Jun Liu and Cornelia van Duijn have used exchange grants from PRECEDI. The funders had no role in study design, data collection and analysis, decision to publish, or preparation of the manuscripts. ERF study is grateful to all study participants and their relatives, general practitioners and neurologists for their contributions and to P. Veraart for her help in genealogy, J. Vergeer for the supervision of the laboratory work and P. Snijders for his help in data collection. The scripts and analysis pipeline used for homozygosity mapping belong to the ROHGEN consortium (https://www.wiki.ed.ac.uk/display/ROHgen/ROHgen2) which ERF is a part of. We thank David Clarke and Jim Wilson for sharing the pipeline.

## Author Contributions

A.D, J.L. and C.M.v.D contributed to study design. K.W.v.D. and C.M.v.D contributed to data collection. A.D., J.L., N.A., and J.B.v.K. contributed to data analysis. A.D., J.L. contributed to writing of manuscript. All the authors contributed to critical review of manuscript.

