## Supplementary for "Recessive genetic effects on type 2 diabetes-related metabolites in a consanguineous population"

**Supplementary Table 1**. Association with exonic SNVs in the loci (NGS based measurements) using the recessive genetic model

| Chr | Start | Stop | Trait | N total/exonic | Bonferroni P-value total/exonic | Minimum P-value total/exonic | RsID SNV/exonic SNV | Gene SNV/exonic SNV |
|---|---|---|---|---|---|---|---|---|
| 2 | 45M | 48M | XL-HDL-cholesterol | 98/12 | $5.10 \times 10^{-4}$ / $4.16 \times 10^{-3}$ | ***$8.94 \times 10^{-5}$/*** $3.24 \times 10^{-2}$ | ***rs57182920*** / rs17480869 | TTC7A |
| 2 | 162M | 165M | Valine*, Isoleucine, Leucine | 43/6 | $1.16 \times 10^{-3}$ / $8.33 \times 10^{-3}$ | $1.50 \times 10^{-2}$ / $6.49 \times 10^{-2}$ | 2:162135276 / rs17783344 | |
| 3 | 63M | 66M | T2DM | 33/2 | $1.52\times 10^{-3}$ / $2.5 \times 10^{-2}$ | $6.99 \times 10^{-3}$ / $4.32 \times 10^{-1}$ | rs66979216 / rs1053338 | |
| 3 | 66M | 69M | T2DM | 34/9 | $1.47\times 10^{-3}$ / $5.55 \times 10^{-3}$ | $4.14 \times 10^{-2}$ / $6.57 \times 10^{-2}$ | rs3828366 / rs6549143 | |
| 3 | 69M | 72M | M-VLDL-triglycerides*, M-VLDL-phospholipids | 36/7 | $1.39 \times 10^{-3}$ / $7.14 \times 10^{-3}$ | ***$1.10 \times 10^{-3}$ / $2.31 \times 10^{-3}$*** | ***rs73095903 / rs62254461*** | FRMD4B / FRMD4B |
| 4 | 27M | 30M | M-LDL-ApoB*, M-LDL-phospholipids, S-LDL-Free cholesterol | 7/0 | $7.14 \times 10^{-3}$ / na | $1.69 \times 10^{-1}$ / na | rs6822297 / na | |
| 4 | 30M | 33M | S-LDL-cholesterol, S-LDL-Free cholesterol*, S-LDL-phospholipids, S-VLDL-triglycerides | 9/0 | $5.56 \times 10^{-3}$ / na | $1.04 \times 10^{-1}$ / na | rs78000988 / na | |
| 4 | 36M | 39M | S-LDL-Free cholesterol, S-LDL-phospholipids, S-LDL-cholesterol, S-LDL-ApoB* | 70/25 | $7.14 \times 10^{-4}$ / $2.00 \times 10^{-3}$ | $1.67\times 10^{-3}$ / $9.82 \times 10^{-3}$ | rs4833106 / rs10776482 | |
| 4 | 39M | 42M | Citrate | 112/25 | $4.46 \times 10^{-4}$ / $2.00 \times 10^{-3}$ | $2.54\times 10^{-2}$ / $8.99 \times 10^{-2}$ | rs4975015 / rs2125313 | |
| 4 | 63M | 66M | Phosphatidylcholine diacyl C 40:6 | 8/0 | $6.25\times 10^{-3}$ / na | ***$7.86 \times 10^{-4}$ /*** na | rs6551721 / na | EXOC5P1 |
| 4 | 66M | 69M | VLDL-ApoB | 74/19 | $6.76\times 10^{-4}$ / $2.63 \times 10^{-3}$ | $1.27 \times 10^{-2}$ / $3.76 \times 10^{-2}$ | rs34321123 / rs1371932 | |
| 6 | 105M | 108M | Phosphatidylcholine diacyl C 40:6 | 39/9 | $1.28 \times 10^{-3}$ / $5.55 \times 10^{-3}$ | $1.11\times 10^{-2}$ / $4.39 \times 10^{-2}$ | rs4946811 / rs9486069 | |
| 6 | 108M | 111M | S-VLDL-triglycerides | 106/16 | $4.72 \times 10^{-4}$ / $3.12 \times 10^{-3}$ | $2.95 \times 10^{-2}$ / $8.56 \times 10^{-2}$ | rs149155788 / rs1040285 | |
| 6 | 111M | 114M | S-LDL-ApoB, S-VLDL-triglycerides* | 41/5 | $1.22 \times 10^{-3}$ / $1.00 \times 10^{-2}$ | $1.98\times 10^{-2}$ / $2.13 \times 10^{-2}$ | rs4947093 / rs33965856 | |
| 8 | 102M | 105M | S-HDL-ApoA1, S-HDL-phospholipids | 76/14 | $6.58 \times 10^{-4}$ / $3.57 \times 10^{-3}$ | $5.67\times 10^{-2}$ / $9.4708 \times 10^{-2}$ | rs56136557 / rs3736043 | |
| 8 | 105M | 108M | S-HDL-ApoA1 | 41/10 | $1.22 \times 10^{-3}$ / $5.00 \times 10^{-3}$ | $3.78\times 10^{-3}$ / ***$3.78 \times 10^{-3}$*** | rs1681904 / ***rs1681904*** | OXR1 |

| | | | | | | | | |
|---|---|---|---|---|---|---|---|---|
| 8 | 114M | 117M | S-HDL-ApoA1 | 12/2 | $4.17 \times 10^{-3}$ / $2.5 \times 10^{-2}$ | ***$3.79\times 10^{-3}$ /*** $3.90 \times 10^{-1}$ | ***rs72685825*** / rs2219898 | CSMD3 |
| 8 | 117M | 120M | S-HDL-ApoA1 | 37/8 | $1.35 \times 10^{-3}$ / $6.25 \times 10^{-3}$ | $5.11\times 10^{-2}$ / $1.14 \times 10^{-1}$ | rs10283129 / rs16888728 | |
| 9 | 15M | 18M | Citrate | 77/10 | $6.49 \times 10^{-4}$ / $5.00 \times 10^{-3}$ | $8.16\times 10^{-2}$ / $2.49 \times 10^{-1}$ | rs62541923 / rs1539172 | |
| 9 | 78M | 81M | IDL-Free cholesterol | 102/42 | $4.90 \times 10^{-4}$ / $1.19 \times 10^{-3}$ | $2.06\times 10^{-2}$ / $5.09 \times 10^{-2}$ | rs2039323 / rs2077181 | |
| 10 | 0M | 3M | M-VLDL-triglycerides*, M-VLDL-phospholipids | 63/8 | $7.94 \times 10^{-4}$ / $6.25 \times 10^{-3}$ | $1.57\times 10^{-2}$ / $5.13 \times 10^{-1}$ | rs10904535 / rs10904083 | |
| 10 | 126M | 129M | WHR | 107/15 | $4.67 \times 10^{-4}$ / $3.33 \times 10^{-3}$ | $1.63\times 10^{-2}$ / $3.79 \times 10^{-2}$ | 10:126302009 / rs41307583 | |
| 12 | 15M | 18M | XL-LDL-phospholipids | 35/3 | $1.43 \times 10^{-3}$ / $1.66 \times 10^{-2}$ | $4.08\times 10^{-3}$ / $1.06\times 10^{-1}$ | rs2300289 / rs3942536 | |
| 17 | 39M | 42M | T2DM | 290/60 | $1.72 \times 10^{-4}$ / $8.33 \times 10^{-4}$ | $1.09\times 10^{-3}$ / $4.801 \times 10^{-3}$ | rs2074162 / rs903 | |
| 20 | 27M | 30M | S-LDL-ApoB, S-LDL-cholesterol, S-LDL-Free cholesterol, S-LDL-phospholipids | 20/9 | $2.50 \times 10^{-3}$ / $5.55 \times 10^{-3}$ | $3.04\times 10^{-3}$ / $5.41 \times 10^{-1}$ | rs41293110 / 20: 29607441 | |
| 20 | 39M | 42M | IDL-phospholipids | 33/1 | $1.52 \times 10^{-3}$ / $5.00 \times 10^{-2}$ | $7.87\times 10^{-3}$ / $1.72\times 10^{-2}$ | 20:40899151 / rs2425516 | |
| 20 | 45M | 48M | IDL-Free cholesterol, IDL-phospholipids | 67/22 | $7.46 \times 10^{-4}$ / $2.27 \times 10^{-3}$ | ***$4.52 \times 10^{-4}$ / $4.52 \times 10^{-4}$*** | ***rs3746820 / rs3746820*** | PREX1 |
| 20 | 60M | 63M | glucose | 186/56 | $2.69 \times 10^{-4}$ / $8.93 \times 10^{-4}$ | ***$2.79 \times 10^{-9}$ /*** $3.12 \times 10^{-3}$ | ***rs35653162*** / rs1048802 | LAMA5 |
| 22 | 45M | 48M | S-HDL-phospholipids | 169/57 | $2.96 \times 10^{-4}$ / $8.77 \times 10^{-4}$ | $1.60\times 10^{-2}$ / $1.20 \times 10^{-1}$ | rs2018279 / rs4508712 | |

**Chr, Start, Stop**: indicate the borders of the locus included in association analyses; **Trait**: metabolomics traits or T2DM and related traits studies as outcomes; **N total/exonic**: Number of SNVs that are found to be homozygous at least among 5 individuals / Number of SNVs that are found to be homozygous at least among 5 individuals and have functional annotation indicating synonymous, non-synonymous, stop codon, UTR and splice variant; **Bonferroni P-value total/exonic**: Regional P-value threshold calculated according to the number of total/exonic SNVs; **Minimum P-value total/exonic**: P-value for the association of the top most significant SNV in the region for all SNVs/ for the exonic SNVs; **RsID total/exonic** of the of the top most significant SNV in the region for all SNVs/ for the exonic SNVs; **Gene total/exonic** annotated to the top SNV/exonic SNV.

**Supplementary Table 2.** Association with exonic SNVs in the loci (chip based measurements) using the recessive genetic model

| Chr | Start | Stop | Trait | N total/exonic | Bonferroni P-value total/exonic | Minimum P-value total/exonic SNVs | SNV/exonic SNV | Gene SNV/exonic SNV |
|---|---|---|---|---|---|---|---|---|
| 2 | 45M | 48M | XL-HDL-cholesterol | 2/0 | $2.50 \times 10^{-2}$ /na | $1.80 \times 10^{-1}$ /na | rs12472172 / na | |
| 2 | 162M | 165M | Valine*, Isoleucine, Leucine | 6/1 | $8.33 \times 10^{-3}$ / $5.00 \times 10^{-2}$ | ***$2.37 \times 10^{-3}$*** /$2.32 \times 10^{-1}$ | rs6759814 / rs17783344 | KCNH7 |
| 3 | 63M | 66M | T2DM | 4/0 | $1.25 \times 10^{-2}$ /na | $5.59 \times 10^{-2}$ /na | rs9860730 / na | |
| 3 | 66M | 69M | T2DM | 9/5 | $5.56 \times 10^{-3}$ /$1.00 \times 10^{-2}$ | $7.98 \times 10^{-2}$ / $1.78 \times 10^{-1}$ | rs1352855 / rs332374 | |
| 3 | 69M | 72M | M-VLDL-triglycerides*, M-VLDL-phospholipids | 3/1 | $1.67 \times 10^{-2}$ /$5.00 \times 10^{-2}$ | $2.50 \times 10^{-1}$ /$2.62\times 10^{-1}$ | rs6806528 / rs4361282 | |
| 4 | 27M | 30M | M-LDL-ApoB*, M-LDL-phospholipids, S-LDL-Free cholesterol | 8/0 | $6.25 \times 10^{-3}$ /na | $5.35 \times 10^{-2}$ /na | rs6817648 / na | |
| 4 | 30M | 33M | S-LDL-cholesterol, S-LDL-Free cholesterol*, S-LDL-phospholipids, S-VLDL-triglycerides | 4/0 | $1.25 \times 10^{-2}$ /na | $2.06 \times 10^{-1}$ /na | rs10025618 / na | |
| 4 | 36M | 39M | S-LDL-Free cholesterol, S-LDL-phospholipids, S-LDL-cholesterol, S-LDL-ApoB* | 14/8 | $3.57 \times 10^{-3}$ /$6.25 \times 10^{-3}$ | $3.67\times 10^{-2}$ /$1.00\times 10^{-1}$ | rs9654132 / rs9654132 | |
| 4 | 39M | 42M | Citrate | 15/6 | $3.33 \times 10^{-3}$ /$8.33 \times 10^{-3}$ | $9.85 \times 10^{-2}$ /$9.84\times 10^{-2}$ | rs2437323 / rs2437323 | |
| 4 | 63M | 66M | Phosphatidylcholine diacyl C 40:6 | 3/0 | $1.67 \times 10^{-2}$ /na | $3.93 \times 10^{-1}$ /na | rs11131511 / na | |
| 4 | 66M | 69M | VLDL-ApoB | 7/3 | $7.14 \times 10^{-3}$ /$1.66 \times 10^{-2}$ | $1.97 \times 10^{-1}$ /$1.97\times 10^{-1}$ | rs1056787 / rs1056787 | |
| 6 | 105M | 108M | Phosphatidylcholine diacyl C 40:6 | 3/3 | $1.67 \times 10^{-2}$ /$1.66 \times 10^{-2}$ | $5.02 \times 10^{-1}$ /$5.02\times 10^{-1}$ | rs1159148 / rs1159148 | |
| 6 | 108M | 111M | S-VLDL-triglycerides | 17/11 | $2.94 \times 10^{-3}$ /$4.54 \times 10^{-3}$ | $1.68 \times 10^{-1}$ /$2.72\times 10^{-1}$ | rs6910666 / rs61741720 | |
| 6 | 111M | 114M | S-LDL-ApoB, S-VLDL-triglycerides* | 6/4 | $8.33 \times 10^{-3}$ /$1.25 \times 10^{-2}$ | $1.61 \times 10^{-1}$ /$2.23\times 10^{-1}$ | rs2148709 / rs143587921 | |
| 8 | 102M | 105M | S-HDL-ApoA1, S-HDL-phospholipids | 7/4 | $7.14 \times 10^{-3}$ /$1.25 \times 10^{-2}$ | $2.20 \times 10^{-1}$ /$2.20\times 10^{-1}$ | rs3134296 / rs3134296 | |

| | | | | | | | | |
|---|---|---|---|---|---|---|---|---|
| 8 | 105M | 108M | S-HDL-ApoA1 | 5/1 | $1.00 \times 10^{-2}$ /$5.00 \times 10^{-2}$ | ***$4.35 \times 10^{-3}$*** /$1.55\times 10^{-1}$ | rs6469084 / rs2920048 | intergenic |
| 8 | 114M | 117M | S-HDL-ApoA1 | 6/1 | $8.33 \times 10^{-3}$ /$5.00 \times 10^{-2}$ | $1.00 \times 10^{-2}$ /$9.351\times 10^{-2}$ | rs1382469 / rs2219898 | |
| 8 | 117M | 120M | S-HDL-ApoA1 | 10/4 | $5.00 \times 10^{-3}$ /$1.25 \times 10^{-2}$ | $1.91 \times 10^{-1}$ /$2.66\times 10^{-1}$ | rs11991695 / rs2073618 | |
| 9 | 15M | 18M | Citrate | 14/3 | $3.57 \times 10^{-3}$ /$1.66 \times 10^{-2}$ | $1.76 \times 10^{-1}$ /$1.24\times 10^{-1}$ | rs1780159 / rs1539172 | |
| 9 | 78M | 81M | IDL-Free cholesterol | 29/11 | $1.72 \times 10^{-3}$ /$4.54 \times 10^{-3}$ | $5.03 \times 10^{-2}$ /**5.03**$\times 10^{-2}$ | rs1110222 / rs1110222 | |
| 10 | 0M | 3M | M-VLDL-triglycerides*, M-VLDL-phospholipids | 11/3 | $4.55 \times 10^{-3}$ /$1.66 \times 10^{-2}$ | $2.13 \times 10^{-1}$ /2.13× 10-1 | rs34407608 / rs34407608 | |
| 10 | 126M | 129M | WHR | 10/5 | $5.00 \times 10^{-3}$ /$1.00 \times 10^{-2}$ | $6.27 \times 10^{-2}$ /$6.26\times 10^{-2}$ | rs11245008 / rs11245008 | |
| 12 | 15M | 18M | XL-LDL-phospholipids | 0/0 | *na/na | $6.18 \times 10^{-1}$ /na | rs4764292 / na | |
| 17 | 39M | 42M | T2DM | 30/27 | $1.67 \times 10^{-3}$ /$1.85 \times 10^{-3}$ | $1.72 \times 10^{-3}$ /***1.70*** $\times 10^{-3}$ | rs1050784 / rs1050784 | KRT15 |
| 20 | 27M | 30M | S-LDL-ApoB, S-LDL-cholesterol, S-LDL-Free cholesterol, S-LDL-phospholipids | 1/0 | $5.00 \times 10^{-2}$ /na | $6.41 \times 10^{-1}$ /na | rs4911274 / na | |
| 20 | 39M | 42M | IDL-phospholipids | 15/6 | $3.33 \times 10^{-3}$ /$8.33 \times 10^{-3}$ | ***$3.21 \times 10^{-3}$ /*** $2.08\times 10^{-1}$ | rs6101991 / rs6142884 | intergenic |
| 20 | 45M | 48M | IDL-Free cholesterol, IDL-phospholipids | 13/5 | $3.85 \times 10^{-3}$ /$1.00 \times 10^{-2}$ | $1.10 \times 10^{-1}$ /$1.39\times 10^{-1}$ | rs6018424 / rs17265513 | |
| 20 | 60M | 63M | glucose | 9/6 | $5.56 \times 10^{-3}$ /$8.33 \times 10^{-3}$ | $2.09 \times 10^{-1}$ / $3.88\times 10^{-1}$ | rs6142884 / rs11553387 | |
| 22 | 45M | 48M | S-HDL-phospholipids | 28/20 | $1.79 \times 10^{-3}$ /$2.50 \times 10^{-3}$ | $3.50 \times 10^{-3}$ / $8.57\times 10^{-2}$ | rs8142147 / rs6007897 | |

**Chr, Start, Stop**: indicate the borders of the locus included in association analyses; **Trait**: metabolomics traits or T2DM and related traits studies as outcomes; **N total/exonic**: Number of SNVs that are found to be homozygous at least among 5 individuals / Number of SNVs that are found to be homozygous at least among 5 individuals and have functional annotation indicating synonymous, non-synonymous, stop codon, UTR and splice variant; **Bonferroni P-value total/exonic**: Regional P-value threshold calculated according to the number of total/exonic SNVs; **Minimum P-value total/exonic**: P-value for the association of the top most significant SNV in the region for all SNVs/ for the exonic SNVs; **RsID total/exonic** of the of the top most significant SNV in the region for all SNVs/ for the exonic SNVs; **Gene total/exonic** annotated to the top SNV/exonic SNV.

**Supplementary Table 3.** Association analyses of imputed SNPs within the candidate ROHs using the recessive genetic model

| Chr | Start | Stop | Trait | SNP | MAF | P-value | FDR | Gene region |
|---|---|---|---|---|---|---|---|---|
| 2 | 45M | 48M | XL-HDL-cholesterol | rs4366955 | 0.44 | $4.63 \times 10^{-3}$ | 0.433 | n.d |
| 2 | 162M | 165M | Valine*, Isoleucine, Leucine | rs115230448 | 0.05 | $1.92 \times 10^{-3}$ | 0.889 | n.d |
| 3 | 63M | 66M | T2DM | rs59997916 | 0.05 | $4.03 \times 10^{-7}$ | ***0.003*** | Intronic MAGI1 |
| 3 | 66M | 69M | T2DM | rs80310002 | 0.05 | $1.56 \times 10^{-4}$ | 0.619 | n.d |
| 3 | 69M | 72M | M-VLDL-triglycerides*, M-VLDL-phospholipids | rs77505126 | 0.06 | $2.83 \times 10^{-4}$ | 0.498 | n.d |
| 4 | 27M | 30M | M-LDL-ApoB*, M-LDL-phospholipids, S-LDL-Free cholesterol | rs1488290 | 0.44 | $1.07 \times 10^{-2}$ | 0.613 | n.d |
| 4 | 30M | 33M | S-LDL-cholesterol, S-LDL-Free cholesterol*, S-LDL-phospholipids, S-VLDL-triglycerides | rs10866392 | 0.14 | $1.60 \times 10^{-5}$ | ***0.039*** | Intronic LINC02506 |
| 4 | 36M | 39M | S-LDL-Free cholesterol, S-LDL-phospholipids, S-LDL-cholesterol, S-LDL-ApoB* | rs73240383 | 0.08 | $1.16 \times 10^{-5}$ | ***0.031*** | Intronic NWD2 |
| 4 | 39M | 42M | Citrate | rs10031048 | 0.06 | $9.40 \times 10^{-4}$ | 0.891 | n.d |
| 4 | 63M | 66M | Phosphatidylcholine diacyl C 40:6 | rs7435864 | 0.38 | $4.16 \times 10^{-4}$ | 0.189 | n.d |
| 4 | 66M | 69M | VLDL-ApoB | rs4422464 | 0.49 | $5.25 \times 10^{-4}$ | 0.923 | n.d |
| 6 | 105M | 108M | Phosphatidylcholine diacyl C 40:6 | rs9386463 | 0.48 | $4.93 \times 10^{-4}$ | 0.222 | n.d |
| 6 | 108M | 111M | S-VLDL-triglycerides | rs71562230 | 0.06 | $1.34 \times 10^{-5}$ | ***0.018*** | Intergenic near SLC22A16 |
| 6 | 111M | 114M | S-LDL-ApoB, S-VLDL-triglycerides* | rs62420300 | 0.05 | $4.81 \times 10^{-5}$ | 0.088 | n.d |
| 8 | 102M | 105M | S-HDL-ApoA1*, S-HDL-phospholipids | rs7000710 | 0.10 | $6.53 \times 10^{-4}$ | 0.946 | n.d |
| 8 | 105M | 108M | S-HDL-ApoA1 | rs1823799 | 0.50 | $6.22 \times 10^{-3}$ | 0.290 | n.d |
| 8 | 114M | 117M | S-HDL-ApoA1 | rs62540596 | 0.07 | $3.65 \times 10^{-4}$ | 0.284 | n.d |
| 8 | 117M | 120M | S-HDL-ApoA1 | rs10955841 | 0.31 | $2.16 \times 10^{-2}$ | 0.875 | n.d |
| 9 | 15M | 18M | Citrate | rs76327891 | 0.23 | $6.76 \times 10^{-4}$ | 0.723 | n.d |
| 9 | 78M | 81M | IDL-Free cholesterol | rs1339249 | 0.45 | $1.69 \times 10^{-3}$ | 0.510 | n.d |
| 10 | 0M | 3M | M-VLDL-triglycerides*, M-VLDL-phospholipids | rs2804098 | 0.04 | $2.44 \times 10^{-4}$ | 0.598 | n.d |

| | | | | | | | | |
|---|---|---|---|---|---|---|---|---|
| 10 | 126M | 129M | WHR | rs1351684 | 0.21 | $4.43 \times 10^{-4}$ | 0.918 | n.d |
| 12 | 15M | 18M | XL-LDL-phospholipids | rs11832997 | 0.39 | $4.96 \times 10^{-3}$ | 0.168 | n.d |
| 17 | 39M | 42M | T2DM | rs17675302 | 0.07 | $1.09 \times 10^{-4}$ | 0.118 | n.d |
| 20 | 27M | 30M | S-LDL-ApoB*, S-LDL-cholesterol, S-LDL-Free cholesterol, S-LDL-phospholipids | rs6058951 | 0.13 | $1.79 \times 10^{-3}$ | 0.272 | n.d |
| 20 | 39M | 42M | IDL-phospholipids | rs1047605 | 0.18 | $4.79 \times 10^{-4}$ | 0.640 | n.d |
| 20 | 45M | 48M | IDL-Free cholesterol, IDL-phospholipids* | rs1573707 | 0.48 | $5.62 \times 10^{-5}$ | ***0.022*** | Intronic PTPRT |
| 20 | 60M | 63M | glucose | rs75320186 | 0.12 | $5.62 \times 10^{-7}$ | ***0.001*** | Upstream gene variant BIRC7 |
| 22 | 45M | 48M | S-HDL-phospholipids | rs131901 | 0.35 | $7.41 \times 10^{-4}$ | 0.187 | n.d |

**Chr, Start, Stop**: indicate the borders of the locus included in association analyses; **Trait**: metabolomics traits or T2DM and related traits studies as outcomes; **SNP**: Top most significant SNP in the locus; **MAF**: Minor allele frequency of the SNP; **P-value**: P-value for the association of the top most significant SNP in the region; **FDR**: Region-wide False Discovery Rate; **Gene region** : Closest gene defined based on Ensembl database look-up; n.d: Not determined.

**Supplementary Table 4.** Replication of selected SNVs in Rotterdam Study using the recessive genetic model

| Chr | Start | Stop | Trait | RsID | A1 | A2 | β | P-value | MAF |
|---|---|---|---|---|---|---|---|---|---|
| 2 | 162M | 165M | Valine*, Isoleucine, Leucine | ***rs6759814*** $_{chip}$ | ***C*** | ***T*** | ***0.01*** | ***$1.41 \times 10^{-3}$*** | 0.08 |
| 3 | 63M | 66M | T2DM | rs59997916 $_{1000G}$ | C | T | 0.04 | $4.49 \times 10^{-2}$ | 0.07 |
| 3 | 69M | 72M | M-VLDL-triglycerides*, M-VLDL-phospholipids | rs62254461 $_{seq}$ | G | A | 0.02 | $1.65 \times 10^{-1}$ | 0.11 |
| 4 | 30M | 33M | S-LDL-cholesterol, S-LDL-Free cholesterol*, S-LDL-phospholipids, S-VLDL-triglycerides | rs10866392 $_{1000G}$ | T | G | -0.00 | $7.85 \times 10^{-1}$ | 0.24 |
| 4 | 36M | 39M | S-LDL-Free cholesterol, S-LDL-phospholipids, S-LDL-cholesterol | rs73240383 $_{1000G}$ | C | T | 0.00 | $1.44 \times 10^{-1}$ | 0.15 |
| 6 | 108M | 111M | S-VLDL-triglycerides | rs71562230 $_{1000G}$ | C | T | 0.01 | $1.80 \times 10^{-1}$ | 0.02 |
| 17 | 39M | 42M | T2DM | rs1050784 $_{chip}$ | C | T | 0.00 | $9.77 \times 10^{-1}$ | 0.33 |
| 20 | 39M | 42M | IDL-phospholipids | rs2425516 $_{seq}$ | A | G | -0.00 | $4.88 \times 10^{-1}$ | 0.39 |
| 20 | 39M | 42M | nmrlipo_IGFCmgdL | rs6101991 $_{chip}$ | G | A | 0.00 | $9.40 \times 10^{-1}$ | 0.10 |
| 20 | 45M | 48M | IDL-Free cholesterol, IDL-phospholipids* | ***rs1573707*** $_{1000G}$ | ***C*** | ***T*** | ***0.01*** | ***$1.99 \times 10^{-3}$*** | 0.45 |
| 20 | 45M | 48M | IDL-Free cholesterol,IDL-phospholipids* | rs3746820 $_{seq}$ | G | A | 0.00 | $4.10 \times 10^{-1}$ | 0.17 |
| 20 | 60M | 63M | glucose | rs75320186 $_{1000G}$ | G | T | 0.03 | $6.30 \times 10^{-1}$ | 0.09 |

**Chr, Start, Stop**: indicate the borders of the locus included in association analyses; **Trait**: metabolomics traits or T2DM and related traits studies as outcomes; **RsID**: Top most significant genetic variant selected for replication; **A1:** effect allele**, A2:** non-effect allele**; β:** Effect estimate per 2 copies of A1 versus the rest of the genotypes**; P-value**: P-value for the β**; MAF**: Minor allele frequency of the genetic variant studied. The associations that pass the replication threshold as defined by 0.05/12 (number of genetic variants tested) are marked in italic/bold; **Chip/Seq/1000G:** indicating the genotyping platform of the variant in discovery and replication phase.

### Supplementary Figure 1. ROH overlap between the neighboring regions

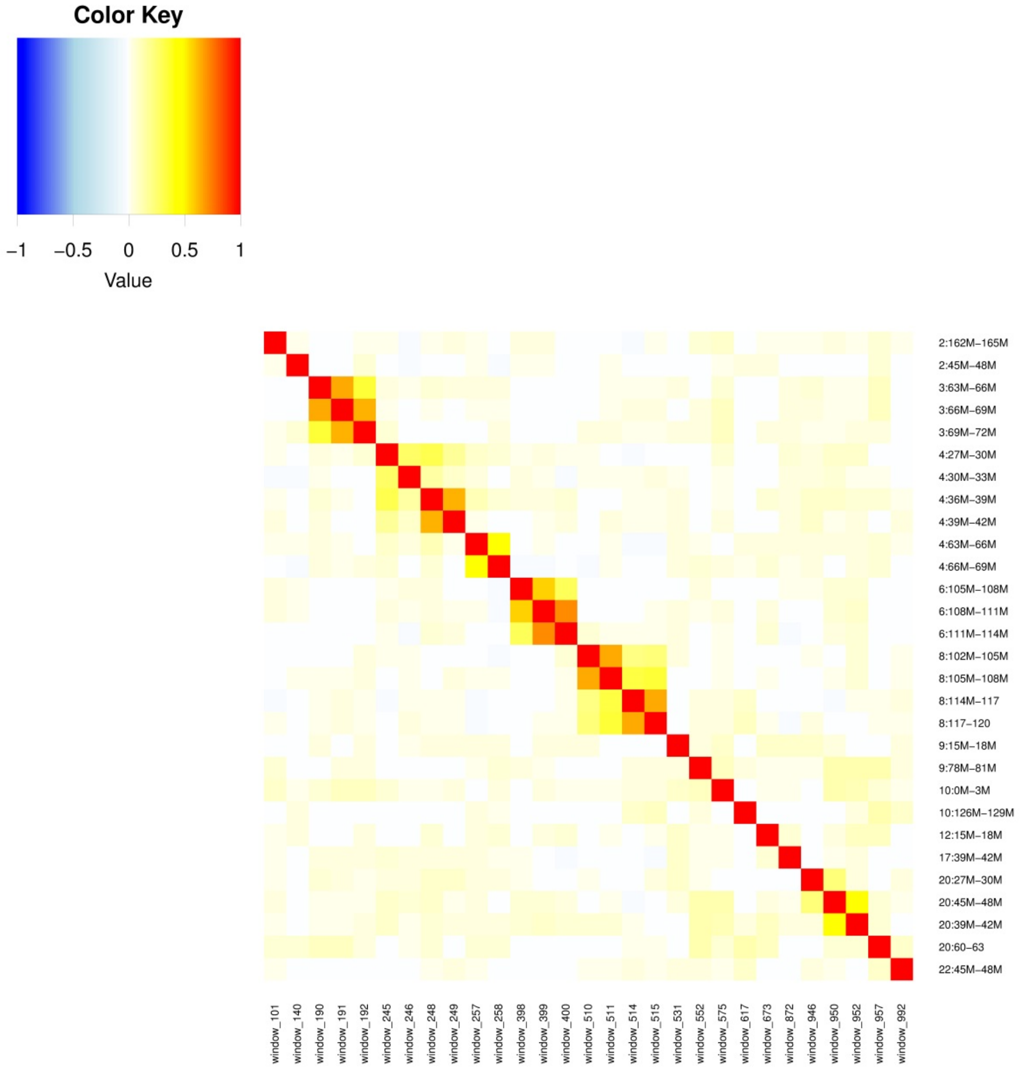

**Supplementary Figure 2. Selected ROH distributions in ERF population**

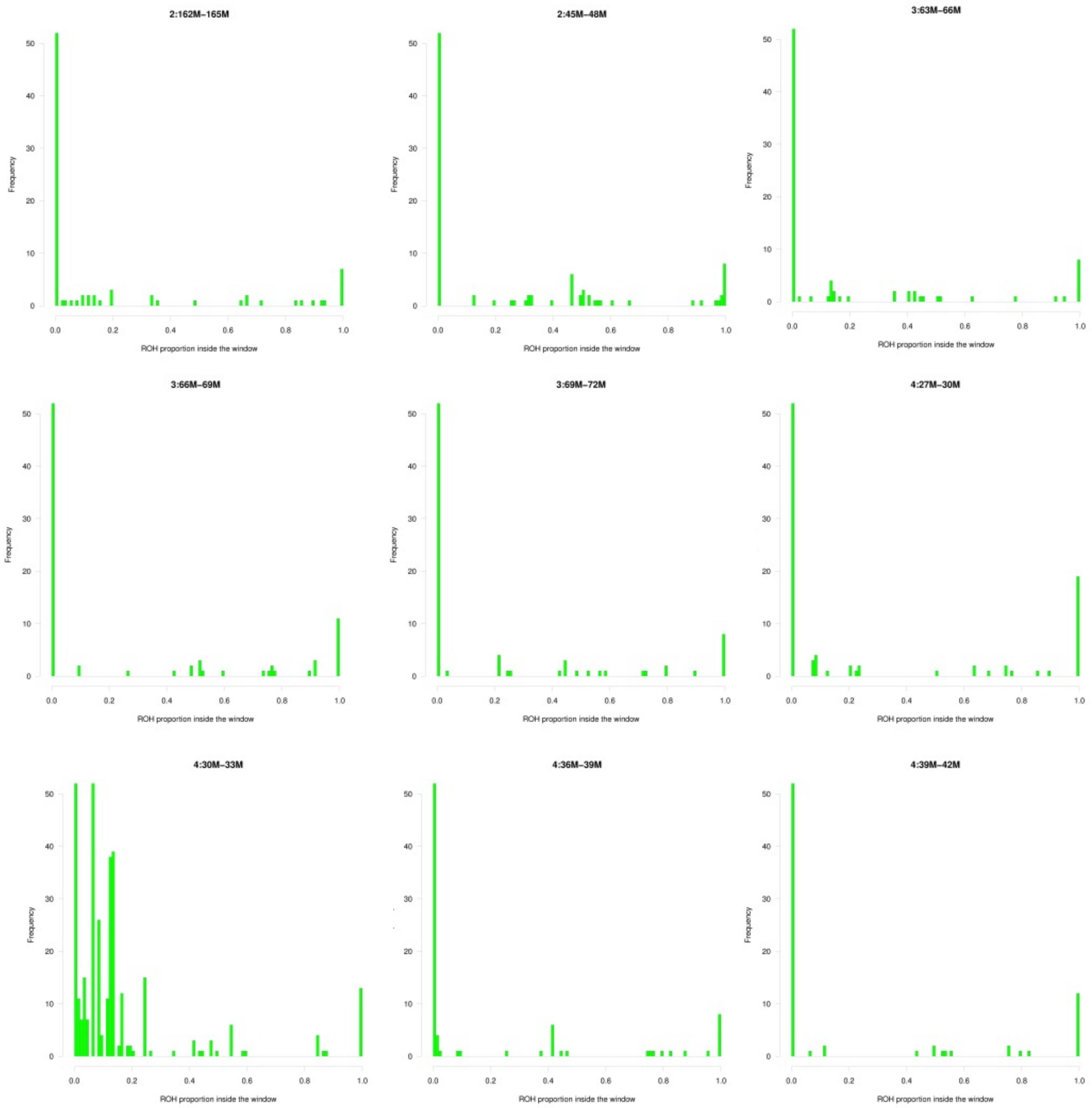

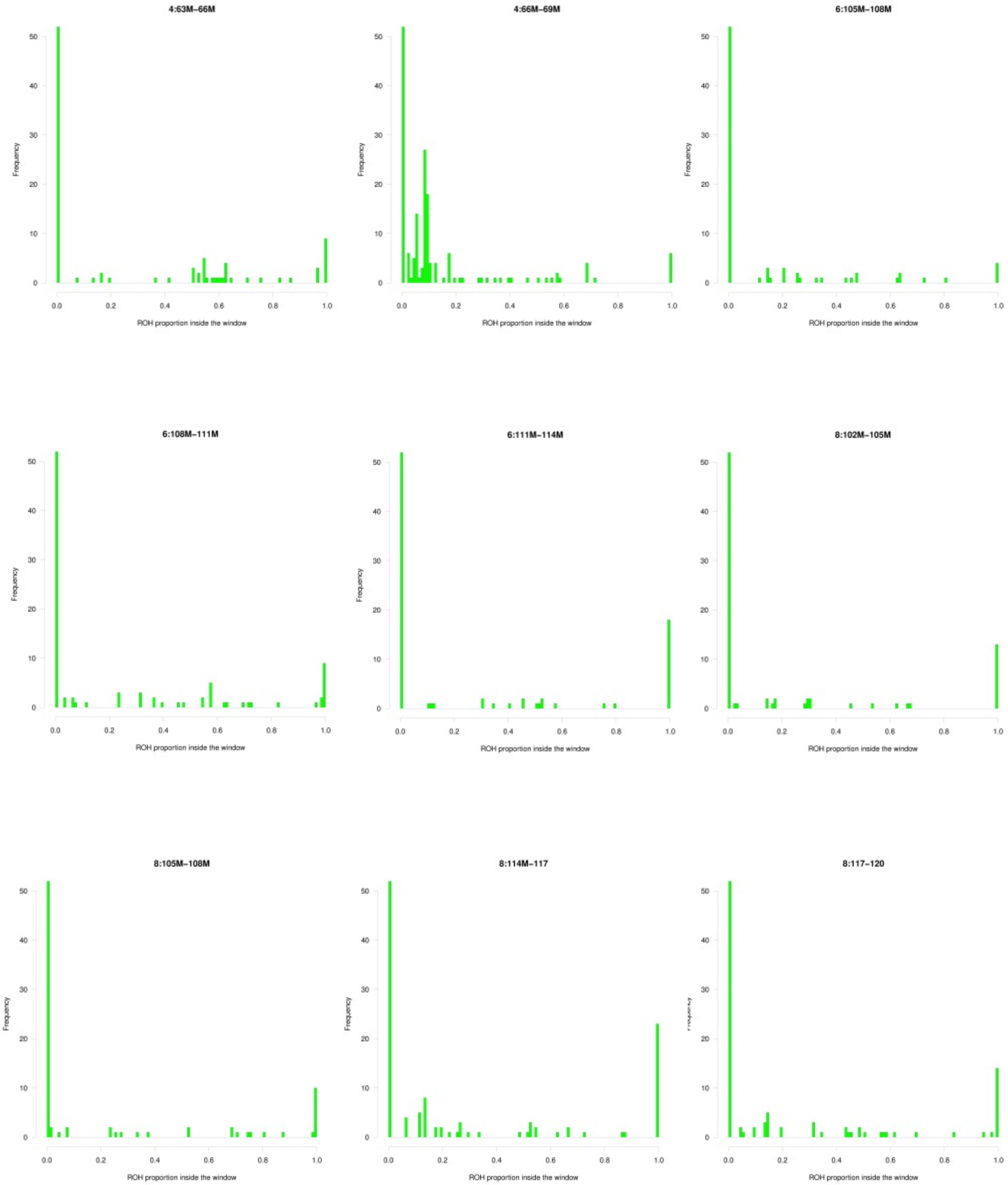
4:63M−66M
4:66M−69M
6:105M−108M
6:108M−111M
6:111M−114M
8:102M−105M
8:105M−108M
8:114M−117
8:117−120
Frequency
ROH proportion inside the window
0.0
0.2
0.4
0.6
0.8
1.0
0
10
20
30
40
50

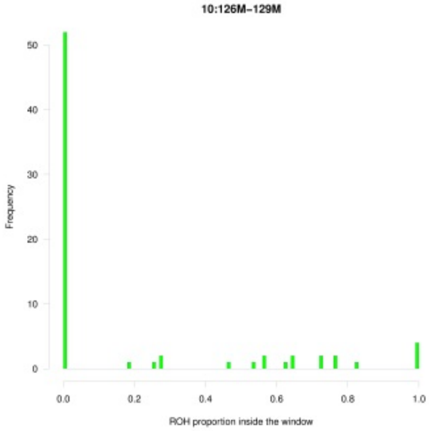
10:126M−129M
Frequency
ROH proportion inside the window

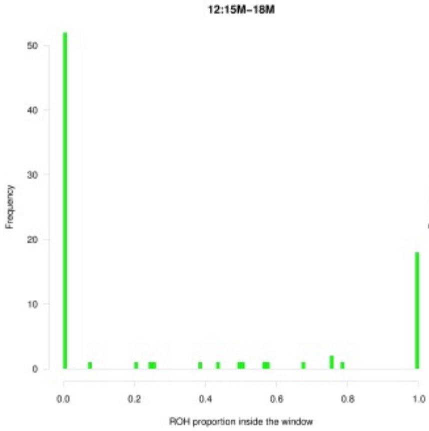
12:15M−18M
Frequency
ROH proportion inside the window

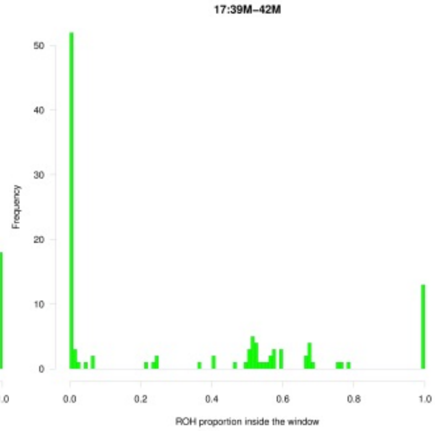
17:39M−42M
Frequency
ROH proportion inside the window

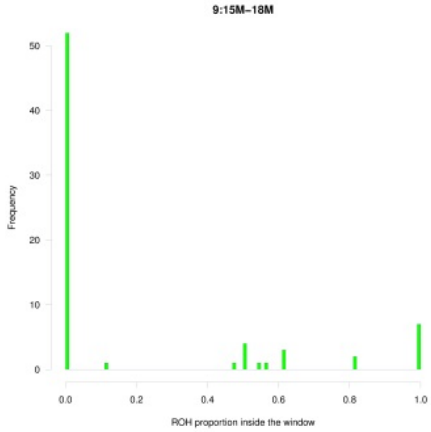
9:15M−18M
Frequency
ROH proportion inside the window

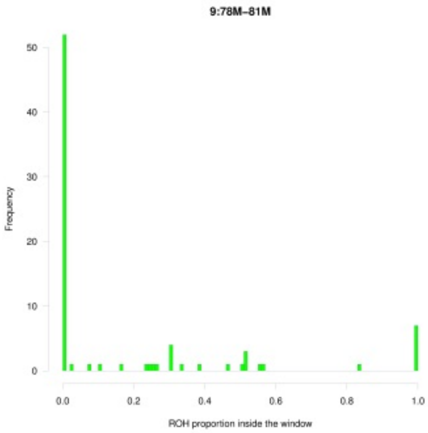
9:78M−81M
Frequency
ROH proportion inside the window

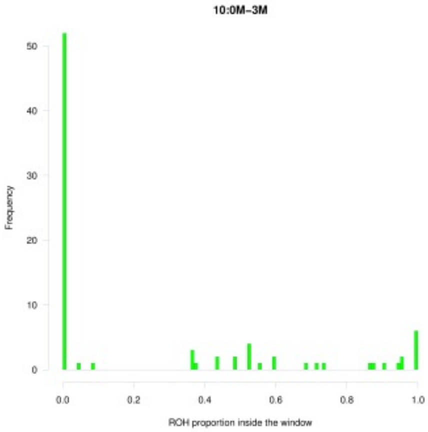
10:0M−3M
Frequency
ROH proportion inside the window

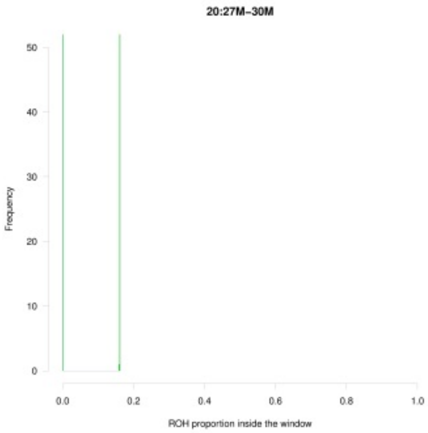
20:27M−30M
Frequency
ROH proportion inside the window

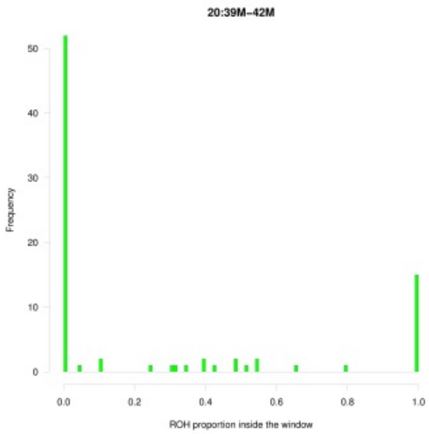
20:39M−42M
Frequency
ROH proportion inside the window

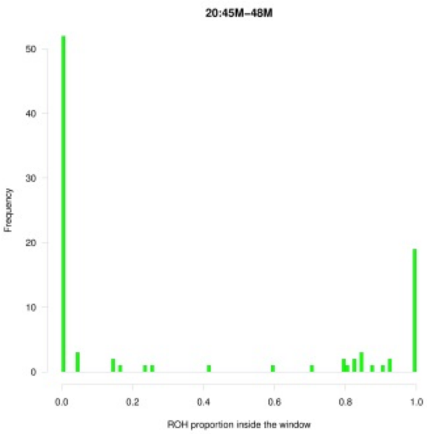
20:45M−48M
Frequency
ROH proportion inside the window

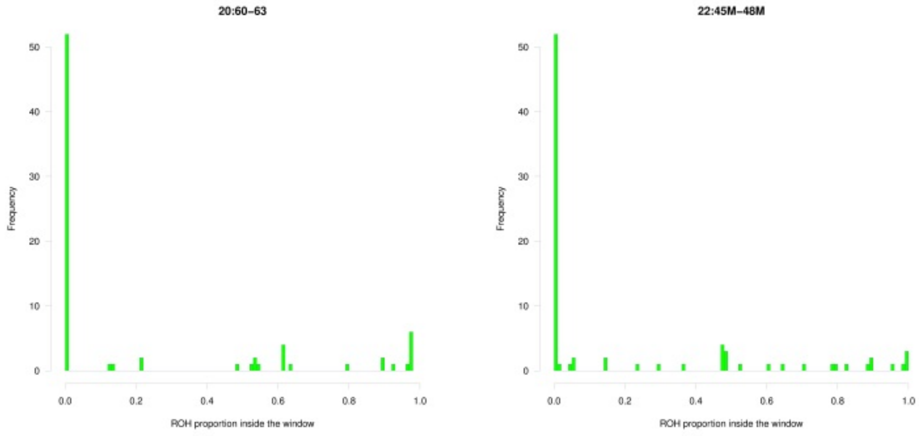

20:60-63
22:45M-48M
Frequency
ROH proportion inside the window

**Supplementary Figure 3a**. Locus zoom plots of association for T2DM under the recessive model within the candidate ROH using 1000G imputed SNPs.

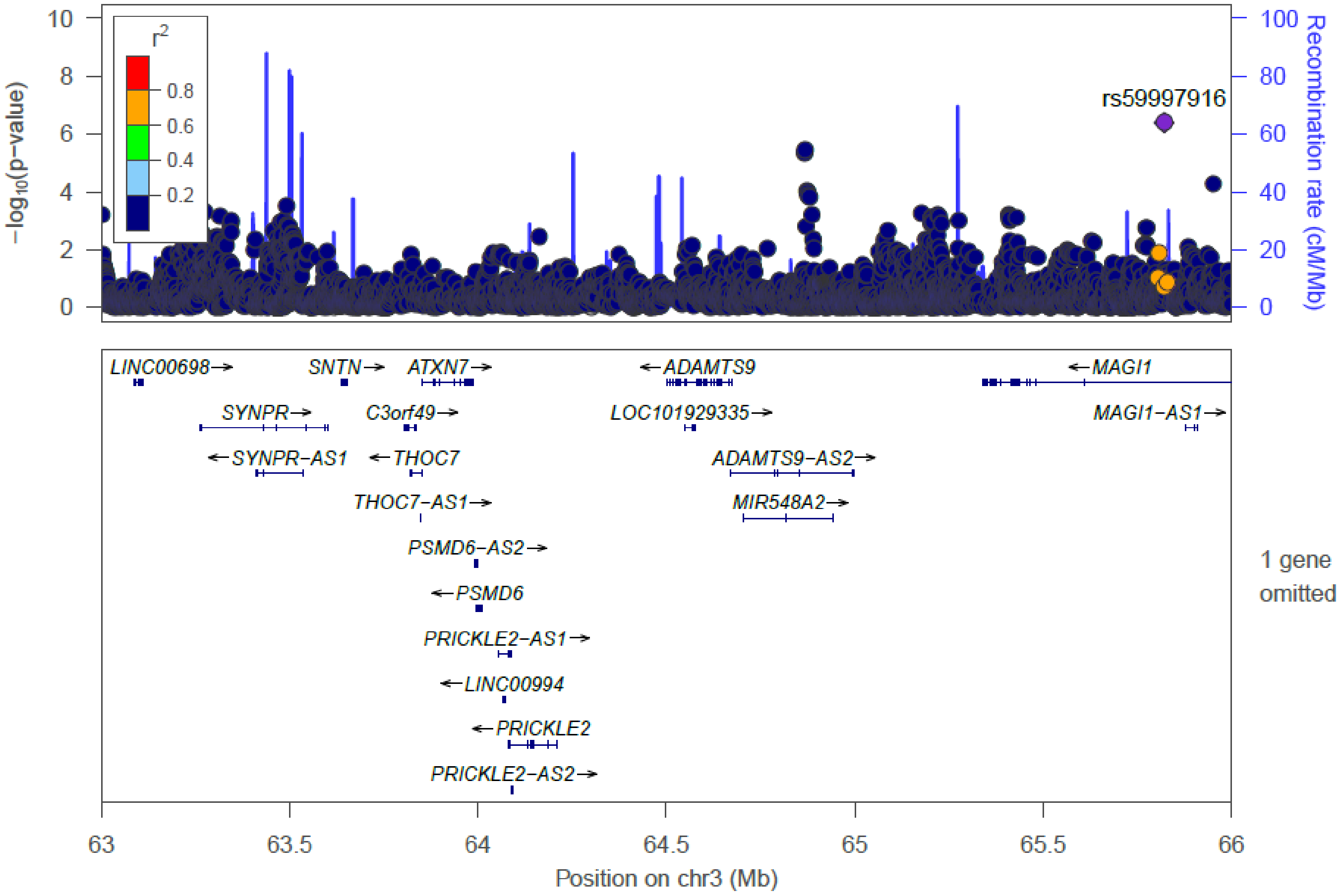

**Supplementary Figure 3b**. Locus zoom plots of association for S-VLDL-TG under the recessive model within the candidate ROH using 1000G imputed SNPs.

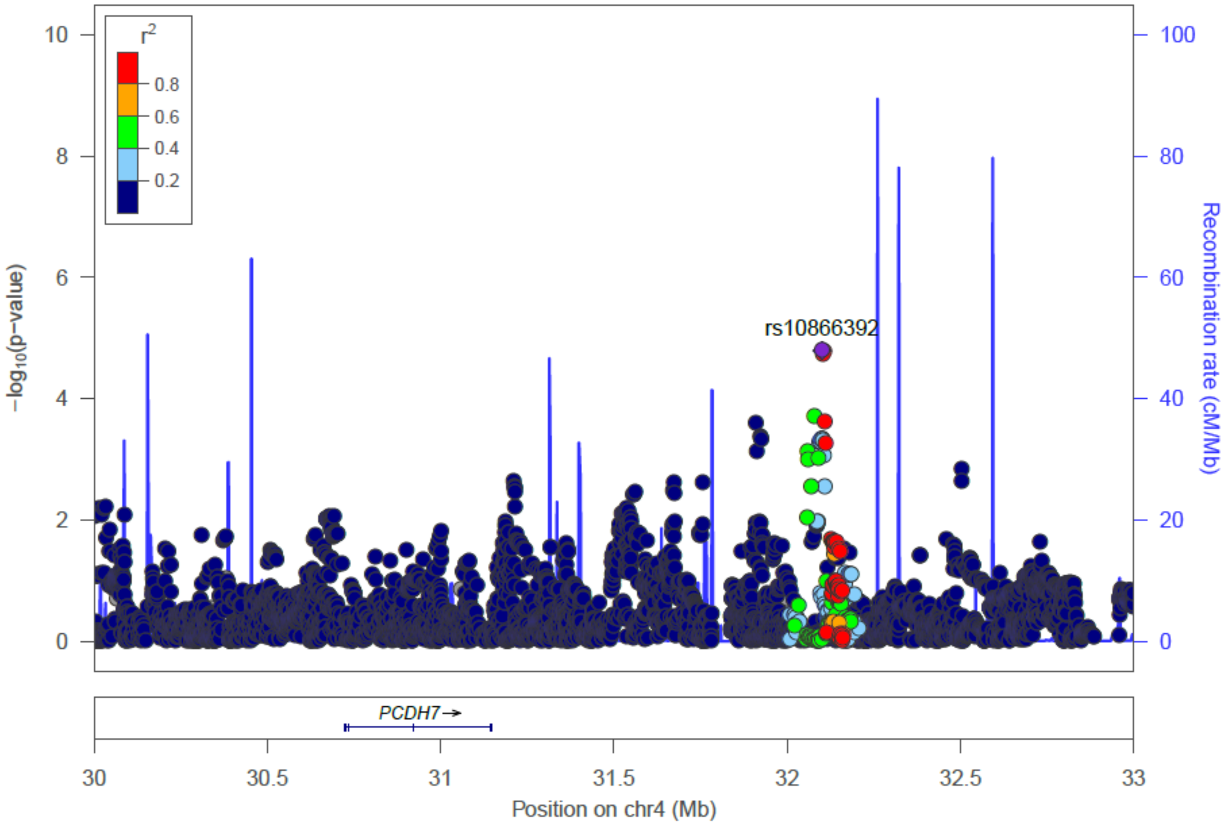

**Supplementary Figure 3c**. Locus zoom plots of association for LDL-Apolipoprotein B under the recessive model within the candidate ROH using 1000G imputed SNPs.

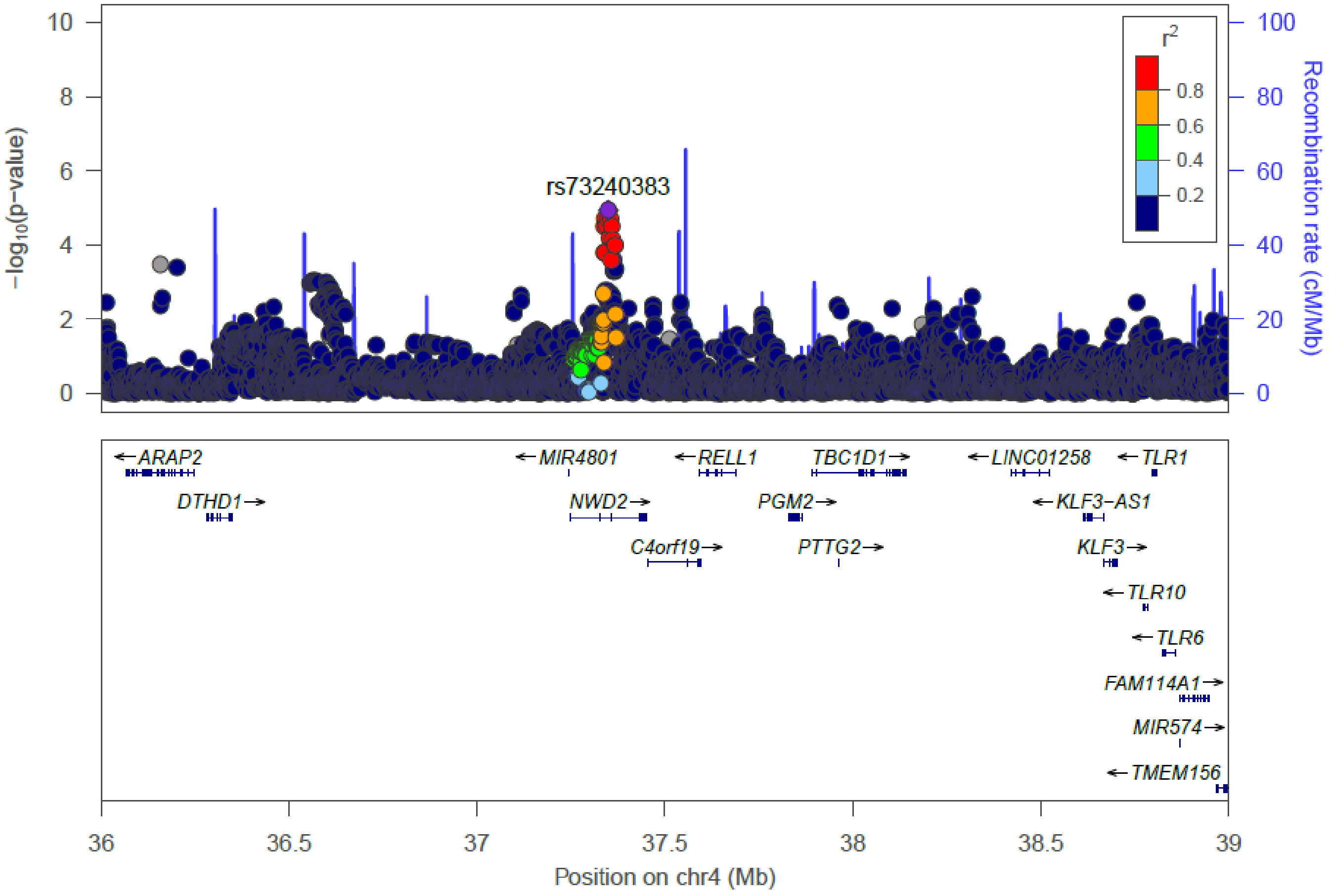

**Supplementary Figure 3d**. Locus zoom plots of association for S-VLDL-TG under the recessive model within the candidate ROH using 1000G imputed SNPs.

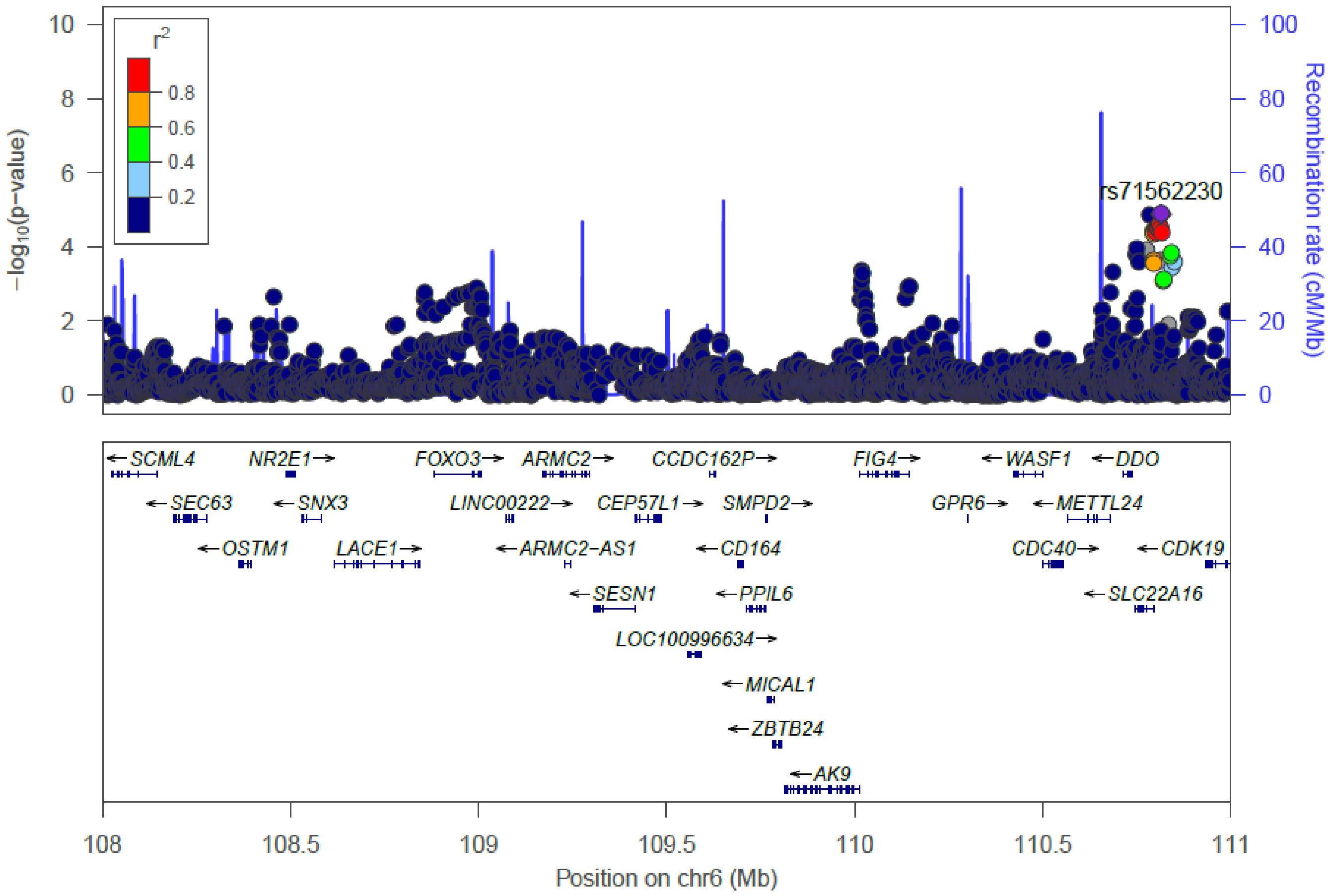

**Supplementary Figure 3e** Locus zoom plots of association for IDL-Free cholesterol under the recessive model within the candidate ROH using 1000G imputed SNPs.

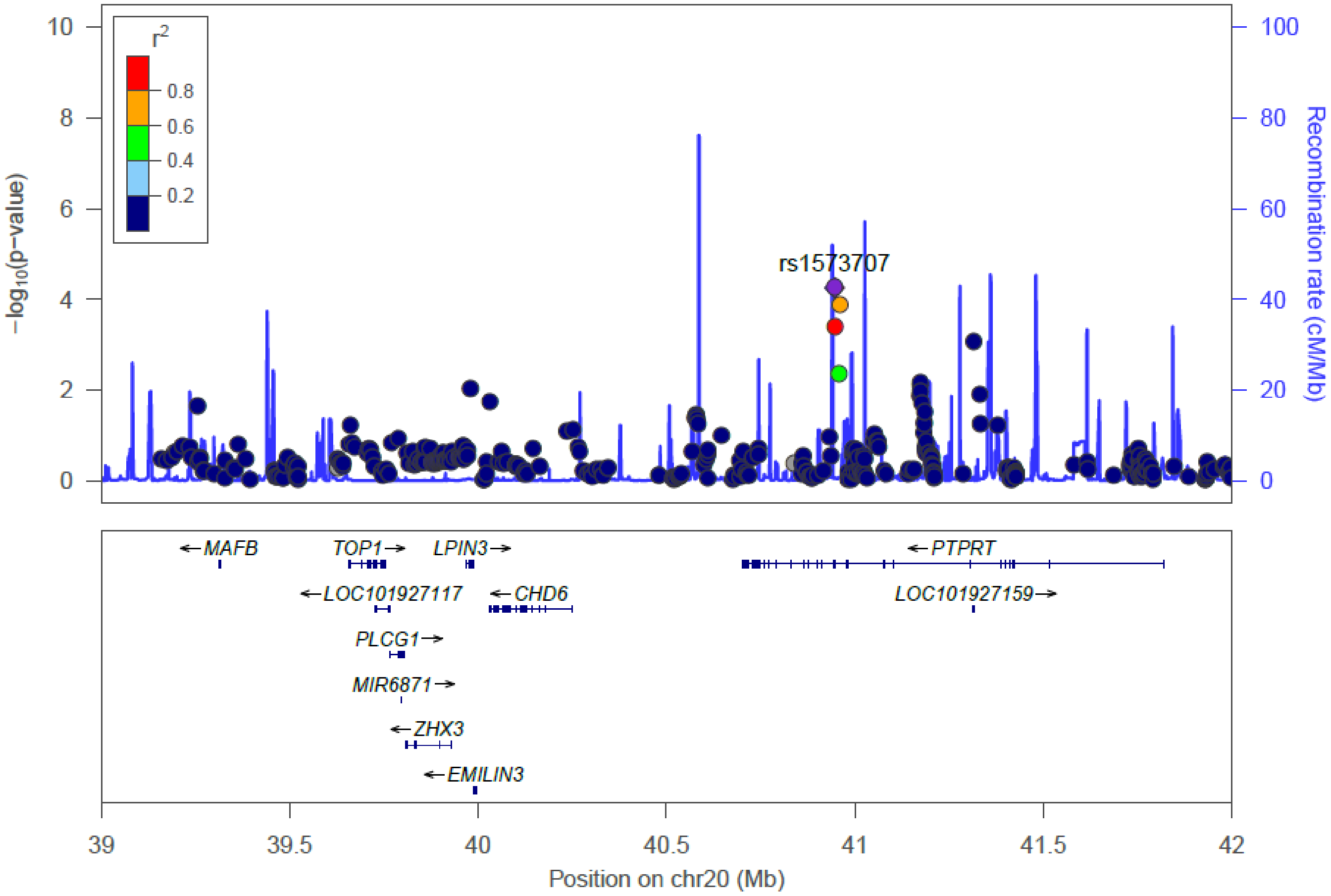

**Supplementary Figure 3f** Locus zoom plots of association for glucose under the recessive model within the candidate ROH using 1000G imputed SNPs.

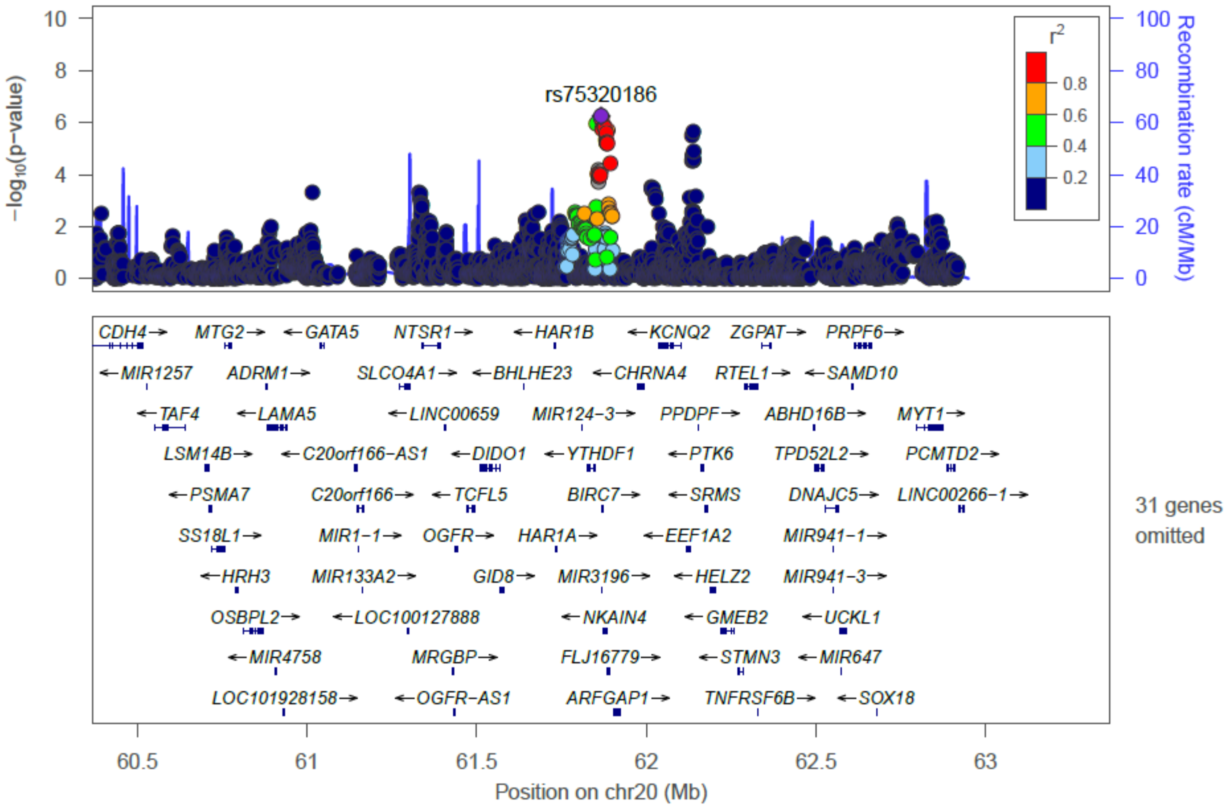
